# NOD1 mediates non-canonical inflammasome processing of interleukin-18 in epithelial cells to *Helicobacter pylori* infection

**DOI:** 10.1101/587212

**Authors:** L. S. Tran, L. Ying, K. D’Costa, G. Wray-McCann, G. Kerr, L. Le, C. C. Allison, J. Ferrand, H. Chaudhry, J. Emery, A. De Paoli, S. Creed, M. Kaparakis-Liaskos, J. Como, J. Dowling, P. A. Johanesen, T. A. Kufer, J. S. Pedersen, A. Mansell, D. J. Philpott, K. Elgass, H. E. Abud, U. Nachbur, B. A. Croker, S. L. Masters, R. L. Ferrero

## Abstract

The interleukin-1 family members, IL-1β and IL-18, are processed into their biologically active forms by multi-protein complexes, known as inflammasomes. Although the inflammasome pathways that mediate IL-1β processing in myeloid cells have been extensively studied, those involved in IL-18 processing, particularly in non-myeloid cells, are still poorly understood. Here, we have identified the cytosolic sensor NOD1 as a key regulator of IL-18 processing in epithelial cells responding to *Helicobacter pylori* infection. Importantly, NOD1 processing of IL-18 occurs independently of the canonical inflammasome proteins, NLRP3 and ASC. Instead, NOD1 interacts directly with caspase-1 via homotypic binding of caspase-activation recruitment domains. We show that IL-18 is important in maintaining tissue homeostasis and protecting against pre-neoplastic changes due to gastric *H. pylori* infection. These findings reveal an unanticipated role for NOD1 in a new type of inflammasome that regulates epithelial cell production of bioactive IL-18 with tissue protective functions.

## Introduction

Interleukin-1 (IL-1) family cytokines (*e.g.* IL-1α, IL-1β, IL-18) play important roles in inflammatory disorders of the gastrointestinal tract ^1^. These cytokines undergo proteolytic cleavage to become biologically active ^1^. The processing of IL-1 family members IL-1β and IL-18 is mediated by multi-protein complexes, known as “inflammasomes”, which can play both beneficial and deleterious roles in the host during infection ^2, 3^. At present, five proteins have been shown to form “canonical inflammasomes”, with several others awaiting confirmation ^2^. The confirmed proteins are: nucleotide-binding domain (NBD) and leucine-rich repeat (LRR)-containing (NLR) protein containing a pyrin domain 1 (NLRP1); NLRP3; NLR caspase activation and recruitment domain-containing protein 4 (NLRC4); absent in melanoma 2 (AIM2); and pyrin. These proteins act as sensors, forming molecular scaffolds for the recruitment of pro-caspase-1, leading to cytokine processing and induction of inflammatory cell death ^2^.

According to the current model of canonical inflammasome formation, sensor proteins with a pyrin domain (PYD) respond to cellular signals by forming oligomeric structures, then bind via PYD–PYD interactions with the adaptor protein, apoptosis-associated speck-like protein (ASC) ^2^. In turn, ASC interacts with pro-caspase-1 through homotypic caspase-activation recruitment domain (CARD) binding, resulting in cleavage of pro-caspase-1 and subsequent processing of IL- 1β/IL-18 into their biologically active forms ^2^. Alternatively, inflammasome sensor proteins may bind directly with caspase-1 via CARD–CARD interactions to initiate cytokine processing (*e.g.* NLRC4) ^2^. These models of canonical inflammasome activation have been developed from studies on IL-1β responses in haematopoietic cells, particularly those of the myeloid cell lineage. In contrast, knowledge regarding inflammasome functions in non-haematopoietic cell lineages, such as epithelial cells, is still poorly understood ^3^. It was suggested that inflammasomes in epithelial cells are more likely to be directed at IL-18 and not IL-1β processing ^3^.

*IL18* is expressed constitutively in the gastrointestinal tract and is upregulated in response to infection by the gastric mucosal pathogen, *Helicobacter pylori* ^4, 5, 6^. The role of IL-18 in gastric *H. pylori* infection is currently unclear, with both pro- ^4, 5, 6^ and anti- ^7^ inflammatory activities having been assigned to this cytokine. *H. pylori* generally causes chronic gastritis, but in approximately 1% of cases, infection leads to gastric cancer ^8^. Gastric carcinogenesis involves a histological progression from chronically inflamed stomach mucosa, to increasingly severe changes in the glandular epithelium, characterised by atrophic gastritis, intestinal metaplasia and dysplasia, resulting finally in invasive carcinoma ^9^.

*H. pylori* strains that harbour a type IV secretion system (T4SS) are more frequently associated with gastric carcinogenesis than strains lacking this factor ^10^. The *H. pylori* T4SS, which is encoded by the cag pathogenicity island (*cag*PAI), is essential for induction of nuclear factor κB (NF-κB)- dependent pro-inflammatory responses in epithelial cells ^11^. It is now clear that *H. pylori* strains harbouring a functional T4SS can activate the NF-κB pathway via at least three mechanisms ^11^. One of these mechanisms involves sensing by the NLR family member nucleotide-binding oligomerisation domain 1 (NOD1), which is highly expressed in gastric epithelial cells and responds to cell wall peptidoglycan fragments (muropeptides) delivered by the T4SS ^12^. Upon activation, NOD1 undergoes self-oligomerisation, leading to the exposure of its CARD and recruitment of the scaffolding kinase protein, receptor-interacting serine-threonine kinase 2 (RIPK2) ^13^. This leads to activation of NF-κB and mitogen-activated protein kinase (MAPK) cascades, resulting in cytokine and chemokine production ^12, 14, 15^.

Here, we identify a new role for NOD1 in the production of bioactive IL-18 by epithelial cells in response to *H. pylori* infection. Furthermore, we show that IL-18 plays a protective role in mouse *H. pylori* infection models and that epithelial cells are the major source of this cytokine in the stomach. Importantly, NOD1-mediated IL-18 processing is independent of the key canonical inflammasome components, NLRP3 and ASC. Instead, NOD1 interacts directly with caspase-1 via CARD–CARD interactions to promote the production of bioactive IL-18. Finally, we show that NOD1 maintains mucosal tissue homeostasis by regulating epithelial cell proliferation and apoptosis. We propose that NOD1-mediated formation of bioactive IL-18 prevents excessive pathology due to chronic *H. pylori* infection.

## Results

### Epithelial cells are a major source of IL-18 in the gastric mucosa

IL-18 is expressed by many different tissues throughout the body, including the gastrointestinal tract, where it is produced by epithelial cells, macrophages and dendritic cells ^1, 16^. In the stomach, IL-18 expression is constitutive, but is upregulated in response to infection by the pathogenic bacterium, *H. pylori* ^4, 5, 6, 7^. Given this observation, together with the fact that epithelial cells are a major target cell population for *H. pylori*–host interactions ^11^, we sought to investigate inflammasome-mediated processing of IL-18 in epithelial cell responses to *H. pylori* infection. It was first necessary, however, to demonstrate epithelial cells as a source of IL-18 in the stomach.

For this, we infected mice with *H. pylori* and then assessed the *Il18* expression levels of the gastric cell populations isolated from these animals by fluorescence-activated cell sorting (FACS). As a comparison, we also measured *Il1b* expression, because it has a very different expression profile in tissues to that of IL-18 ^16^. Consistent with studies in mice and humans ^4, 5, 6, 7^, we detected constitutive IL-18 production in mice, whereas its production was significantly increased in animals at both 1 and 8 weeks post-infection (p.i.), when compared with control animals given Brain Heart Infusion (BHI) broth alone (Fig. 1a; *p* = 0.03 and *p* = 0.004, respectively). In contrast, gastric IL-1β production was only increased at 8 weeks p.i. (Fig. 1b; *p* = 0.05). FACS analyses of isolated gastric mucosal cells from *H. pylori*-infected mice demonstrated that *Il18* gene expression levels in gastric epithelial cells (EpCAM^+^ CD45^−^) were 22-fold higher than those of the immune cell (EpCAM^−^ CD45^+^) population, whereas *Il1b* was highly expressed in immune cells but undetectable in epithelial cells (Fig. 1c). Immunofluorescence analysis further confirmed that EpCAM^+^ cells represent an abundant source of IL-18 in the gastric mucosa (Fig. 1d). Hence, although both IL-18 and IL-1β production is induced in the gastric mucosa during *H. pylori* infection, these cytokines display distinct cellular sources of origin, with IL-18 predominantly originating from epithelial cells.

**Fig. 1.**
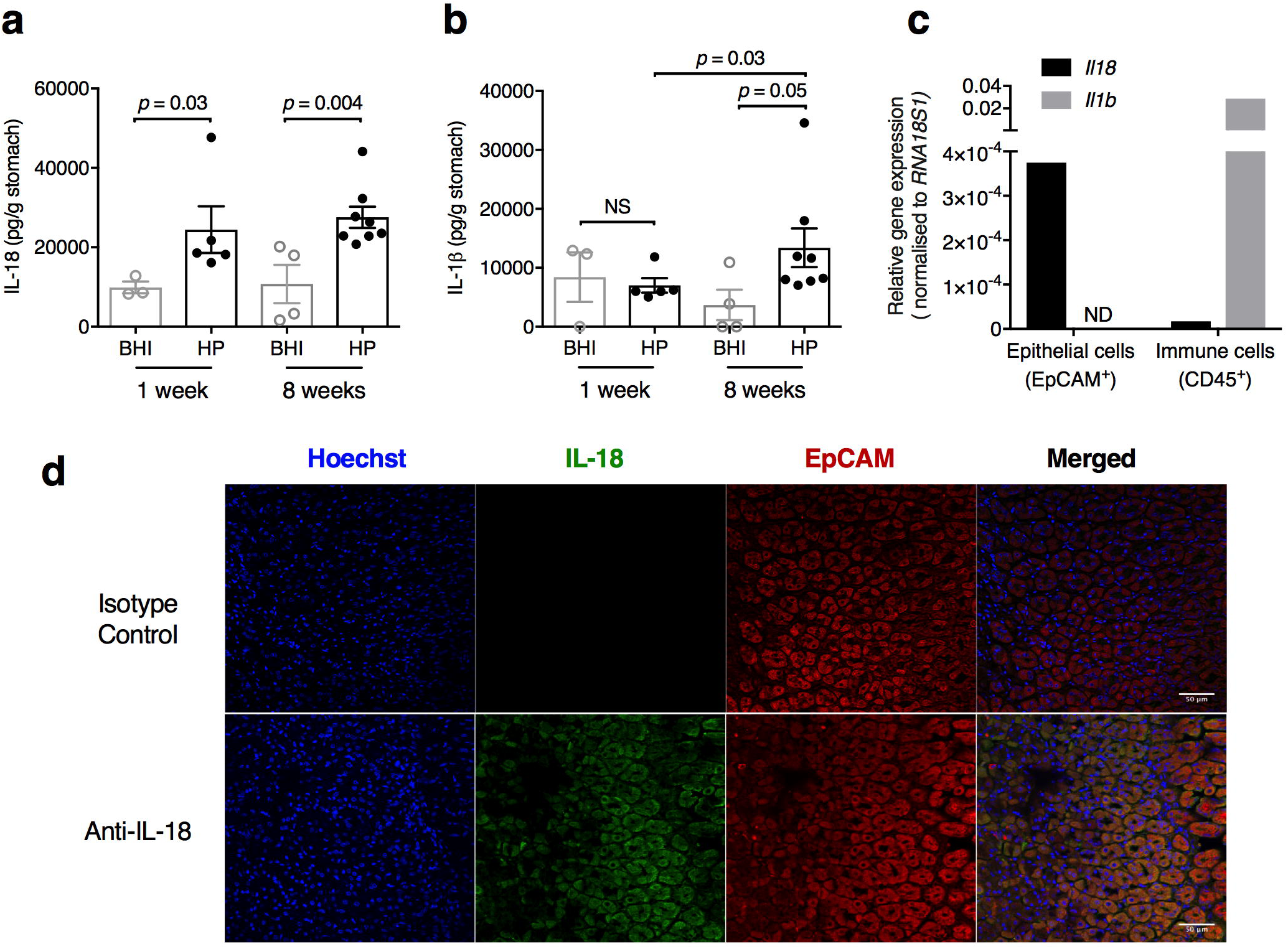
Epithelial cells are a major source of gastric IL-18 in response to *H. pylori* infection. **a** IL-18 and **b** IL-β levels in stomach homogenates from mice administered either BHI broth or *H. pylori* at 1 and 8 weeks p.i. **c** Epithelial (EpCAM^+^ CD45^−^) and immune (CD45.2^+^ EpCAM^−^) cells were isolated by FACS from the gastric tissues of *H. pylori*-infected mice and tested for *Il18* and *Il1b* expression by qPCR. **d** Gastric tissues sections from an *H. pylori*-infected mouse were reacted with rat anti-mouse IL-18 and rabbit anti-EpCAM antibodies, followed by anti-rat Alexa Fluor® 488- and anti-rabbit Alexa Fluor® 594 conjugated secondary antibodies. Mouse IgG was used as an isotype control for IL-18 staining. Cell nuclei were stained with Hoechst 33342 and the sections analysed by confocal microscopy. Scale bar = 50 µm. **a, b** Data pooled from two independent experiments. Significance was determined by the unpaired two-tailed t-test. Data correspond to the mean ± SEM.

### Non-haematopoietic cells produce IL-18 which protects against pre-neoplastic lesions during chronic *H. pylori* infection

To further confirm the epithelial cell origin of IL-18 and also assess its role in *H. pylori* infection, we employed a bone marrow (BM) reconstitution mouse model. Consistent with the idea that epithelial cells are a major source of gastric IL-18, γ-irradiated *Il18*^−/−^ recipient mice that were reconstituted with BM cells either from *Il18*^+/+^ or *Il18*^−/−^ donor mice produced significantly lower levels of IL-18 than *Il18*^+/+^ recipient mice, after 5 weeks p.i. with *H. pylori* (Fig. 2a; *p* = 0.002). Similar findings were observed at 18 weeks p.i. (data not shown). Comparable levels of IL-1β and bacterial loads were observed in both *Il18*^+/+^ and *Il18^−/−^*recipient mice, suggesting that epithelial cell-derived IL-18 does not affect IL-1β production nor bacterial colonisation (Fig. 2b, 2c). Importantly, mice specifically lacking IL-18 in the non-haematopoietic compartment exhibited increased stomach weights (Fig. 2d), mucosal thickness (gastric hyperplasia; Fig. 2d, e) and acid mucin production (Fig. 2f) when compared with animals producing IL-18 in this compartment. Consistent with the findings of the BM reconstitution experiments, we also observed that *Il18*^−/−^ mice exhibit significantly increased stomach weights at 2 months p.i. with *H. pylori*, as compared with *Il18*^+/+^ animals (Supplementary Fig. 1a, 1b), even though the expression levels of inflammatory cytokine genes, *Il1b* and *Cxcl2*, were similar in both groups of animals Supplementary Fig. 1c). Similar differences in stomach weights between *Il18*^+/+^ and *Il18*^−/−^ animals were observed at 13 months p.i. for the related zoonotic *Helicobacter* sp., *Helicobacter felis*, which induces greater inflammatory responses than *H. pylori* in mice ^17^ (Supplementary Fig. 1d, 1e). Taken together, our data show that non-haematopoietic cells in the stomach produce IL-18 which plays a protective role against the development of pre-neoplastic lesions during chronic *Helicobacter* infection.

**Fig. 2.**
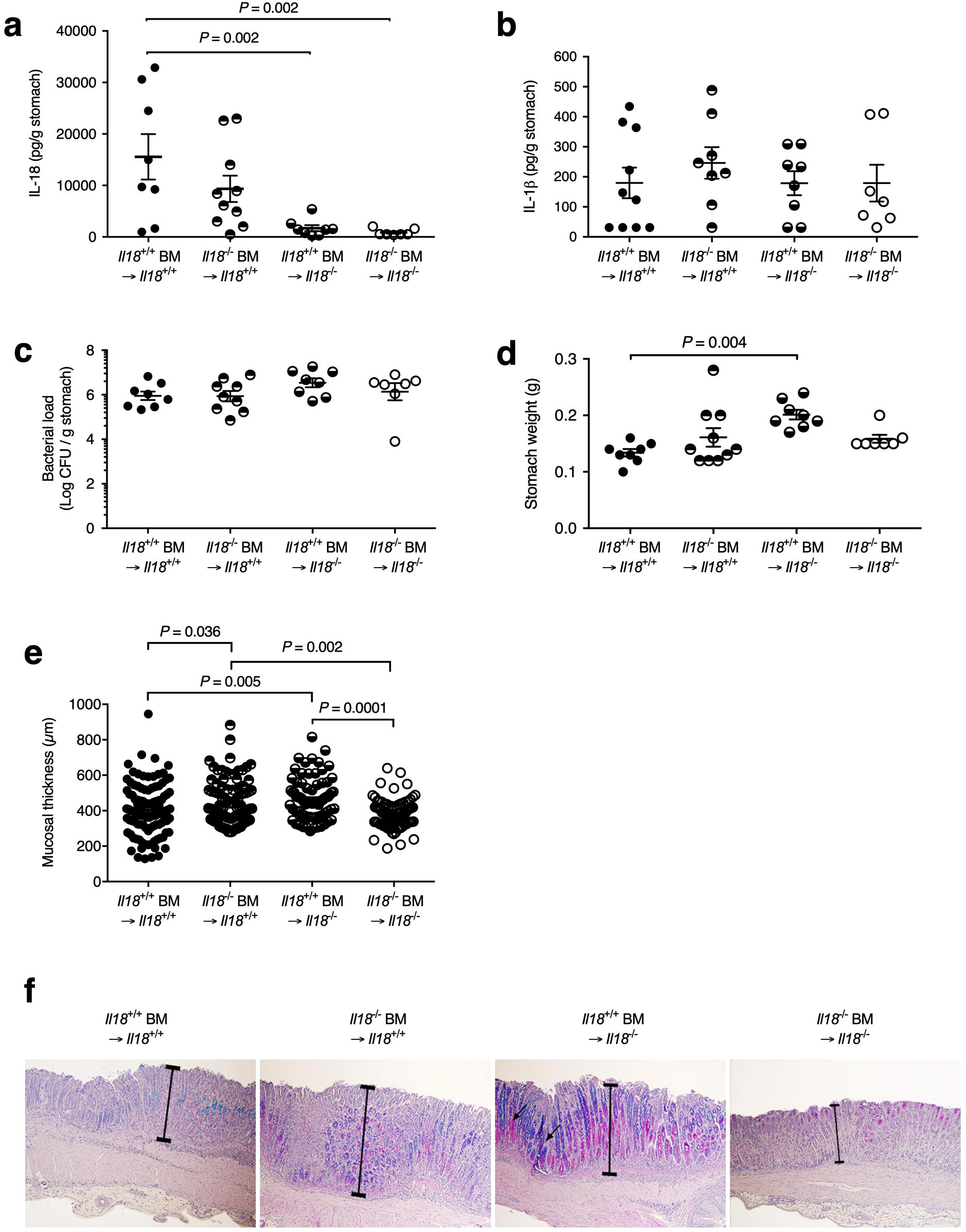
Non-haematopoietic cells produce IL-18 which protects against pre-neoplastic lesions in *H. pylori* infection. BM reconstitution experiments were performed by transferring BM from either *Il18*^+/+^ or *Il18*^−/−^ donor mice to γ-irradiated *Il18*^+/+^ or *Il18*^−/−^ recipient mice which were challenged with *H. pylori* then culled at 5 weeks p.i. **a** IL-18 and **b** IL-β levels in stomach homogenates; **c** stomach *H. pylori* bacterial loads and **d** weights; **e-f** mucosal thickness measurements (e) and images of PAS-AB-stained gastric tissue sections (f). Arrows show acid mucins (blue). **a**-**e** Data pooled from two independent experiments. Individual points refer to either individual mice (**a-d**) or individual data points analysed from 2-4 sections per stomach sample (**e**). Significance was determined by the unpaired two-tailed t-test. Data correspond to the mean ± SEM.

### IL-18 responses to *H. pylori* infection are independent of the canonical inflammasome

Having demonstrated gastric epithelial cells to be a major source of IL-18 in the stomach, we next sought to determine the inflammasome signalling molecules required for its production in response to *H. pylori* infection. For this, we added *H. pylori* bacteria to primary gastric epithelial cells that had been isolated from mice deficient in the canonical inflammasome molecules, Nlrp3, Asc (encoded by the *Pycard* gene) and caspase-1. IL-18 production was significantly reduced in primary cells from *Casp1*^−/−^, but not those from *Nlrp3^−/−^*nor *Pycard^−/−^* mice, when compared with wild-type (WT) cells (Fig. 3a; *p* = 0.004). In contrast to the data in primary epithelial cells, BM-derived macrophages (BMDMs) produced little to no IL-18 in response to stimulation with *H. pylori* when compared with untreated control cells (Supplementary Fig. 2).

**Fig. 3.**
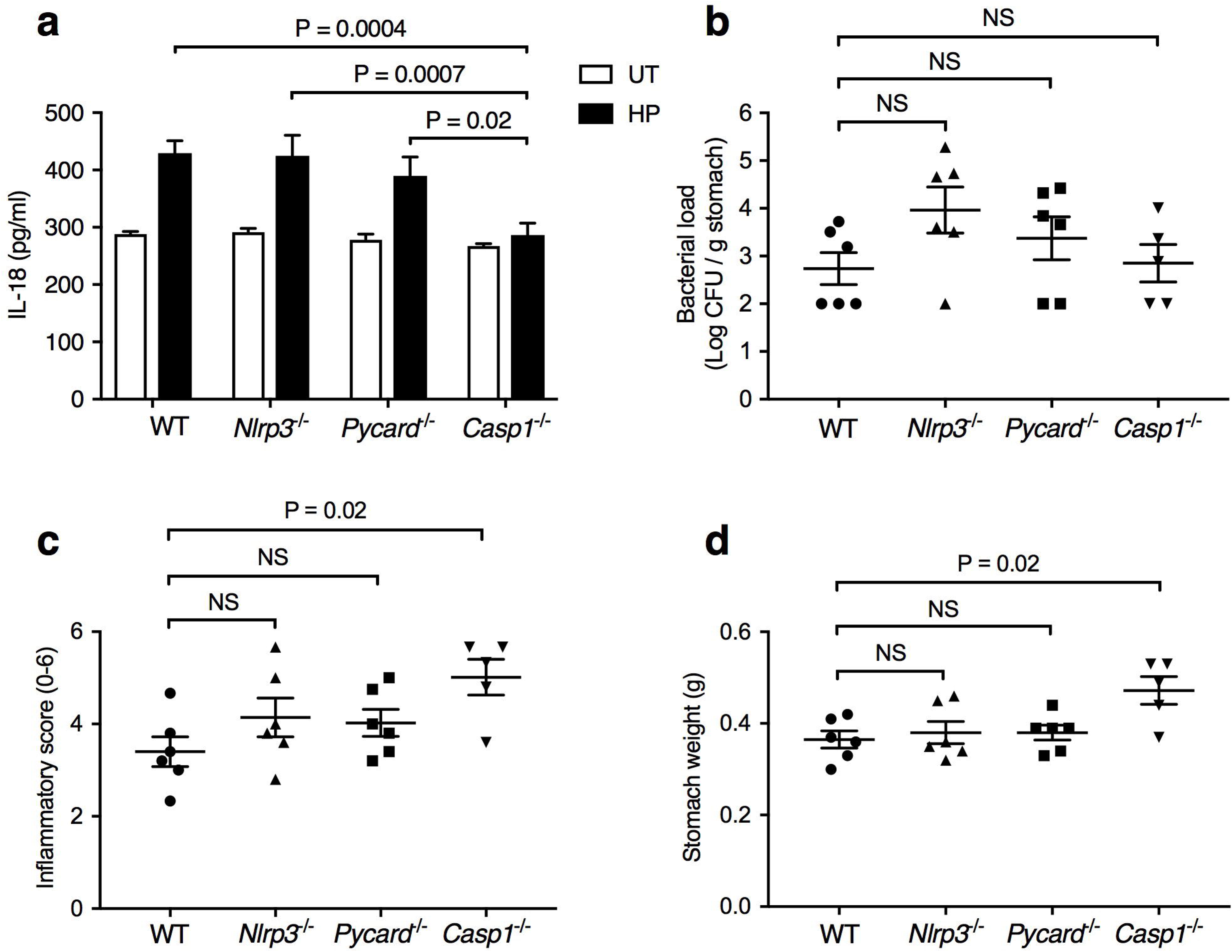
*H. pylori* induces IL-18 production independently of the canonical inflammasome. **a** IL-18 production was assessed in primary gastric epithelial cells that had been isolated from WT, *Nlrp3^−/−^*, *Pycard^−/−^* and *Casp1^−/−^* mice and stimulated with *H. pylori* (HP) bacteria or left untreated (UT). **b-d** WT and KO mouse strains were infected with *H. pylori*, sacrificed at 6 months p.i. and then their stomachs analysed for (**b**) bacterial loads, inflammatory scores (**c**) and weights (**d**). Representative data for three independent experiments (**a**). Significance was determined by the Mann-Whitney test. Data correspond to the mean ± SEM.

Next, we performed *H. pylori* infection studies in WT, *Nlrp3^−/−^*, *Pycard^−/−^* and *Casp1*^−/−^ mice. Although all animal groups showed similar levels of bacterial loads (Fig. 3b), significantly increased inflammatory scores and stomach weights were only observed in *Casp1*^−/−^ mice when compared with WT animals (Fig. 3c, 3d; *p* = 0.02). The increased stomach weights observed in the *Casp1*^−/−^ mice are consistent with the findings for *Il18^−/−^*mice (Fig. 2d, e; Supplementary Fig. 1). Moreover, we infected mice deficient in other known canonical inflammasome molecules, Nlrp1 and Nlrc4, the latter responding to bacterial flagellin and rod proteins ^2^, but observed no significant differences in colonisation levels nor stomach pathology when compared with WT animals (Supplementary Fig. 3). Taken together, the findings suggest that the regulation and functions of IL- 18 in gastric epithelial cells in response to *H. pylori* infection is dependent on caspase-1, but independent of canonical inflammasome pathways, including the key inflammasome molecules NLRP3 and ASC, which have been implicated in responses to bacterial pathogens.

### NOD1 mediates IL-18 processing in response to *H. pylori*

Given the observed role for caspase-1 in *H. pylori*-induced IL-18 processing, we next explored potential caspase-1-interacting proteins that may mediate these responses in epithelial cells. For this, we used the human AGS gastric epithelial cell line, as *H. pylori* stimulation was shown to cause the upregulation of *IL18* mRNA ^6^ and protein ^18^ in these cells. Consistent with the data above, we were unable to detect the presence in AGS cells of either NLRP3 or ASC, whereas both were present in HeLa epithelial and THP-1 monocyte cell lines (Supplementary Fig. 4), suggesting that a non-classical inflammasome molecule may be involved in *H. pylori*-induced IL-18 processing.

In common with primary gastric epithelial cells, AGS cells express the NLR protein NOD1, which plays important roles in the sensing of *H. pylori* bacteria ^12, 14, 15^. Although early studies reported potential CARD–CARD interactions between NOD1 and pro-caspase-1 ^19^, these findings have never been re-visited. To study the potential role of NOD1 in IL-18 processing, we pre-treated cells of the GSM06 mouse gastric epithelial cell line ^20^ with the NOD1-specific inhibitor ML130 ^21^ prior to stimulation with *H. pylori*. Total levels of IL-18 production were increased in the culture supernatants of GSM06 cells in response to *H. pylori* stimulation, whereas it was significantly reduced when NOD1 self-oligomerisation was inhibited by ML130 treatment (Fig. 4a; *p* = 0.03). More significantly, ML130 treatment resulted in reduced production of both precursor and mature forms of IL-18 in these cells (Fig. 4b). These findings were confirmed using primary gastric epithelial cells from *Nod1*^+/+^ and *Nod1^−/−^* mice, in which significantly lower levels of total IL-18 (Fig. 4c; *p* = 0.04), as well as full-length and mature IL-18, were consistently observed in *Nod1^−/−^* gastric epithelial cells as compared to WT cells (Fig. 4c, 4d). Interestingly, BMDMs from *Nod1*^−/−^ mice produced comparable levels of IL-18 (and IL-1β) in response to *H. pylori* and the canonical inflammasome activators, LPS and nigericin (Supplementary Fig. 5), suggesting that NOD1 mediates IL-18 processing in a cell-type-specific manner.

**Fig. 4.**
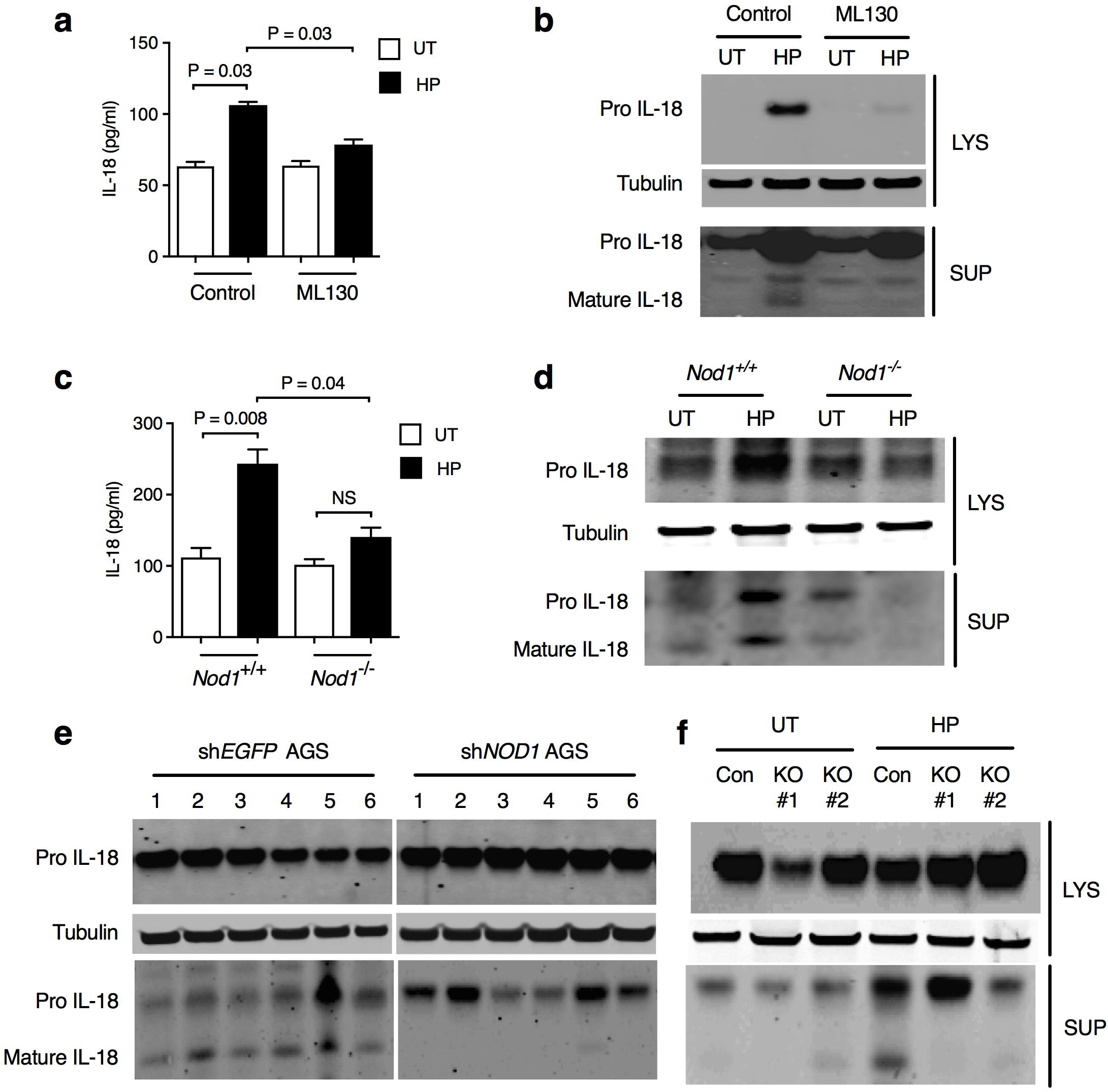
NOD1 is required for *H. pylori*-induced IL-18 processing in epithelial cells. **a, b** Mouse GSM06 gastric epithelial cells that had been pre-treated with NOD1 inhibitor (ML130; 5 µM) or vehicle (control) were stimulated with *H. pylori* (HP) or left untreated (UT). IL-18 levels were measured in culture supernatants (**a**), while pro-IL-18 and mature IL-18 were detected in cell lysates (LYS) and supernatants (SUP), respectively (**b**). Tubulin was used as a loading control. **c, d** Primary gastric epithelial cells from *Nod1*^+/+^ and *Nod1*^−/−^ mice were analysed for IL-18 production (**c**), pro-IL-18 and mature IL-18 (**d**). **e** Human AGS gastric epithelial cells stably expressing shRNA to either *EGFP* (sh*EGFP*) or *NOD1* (sh*NOD1*) were left untreated (**1**), or stimulated with *H. pylori* 251WT (**2**), Δ*cag*PAI (**3**), *cagM* (**4**), 10700 WT (**5**) or SS1 WT (**6**) bacteria. **f** AGS *NOD1* CRISPR/Cas9 KO (two clones) or Cas9 control (CON) cells were stimulated with *H. pylori* (HP) or left untreated (UT). Representative data for three independent experiments (**a-f**). Significance was determined by the unpaired two-tailed t-test (**a**, **c**). NS = not significant. Data correspond to the mean ± SEM.

The contribution of NOD1 to IL-18 processing was further investigated in the AGS human gastric epithelial cell line. Interestingly, interfering with NOD1 signalling in AGS cells by either short hairpin RNA (shRNA) knockdown (sh*NOD1*; ^14, 22^) or CRISPR/Cas9 knockout (*NOD1* KO; 15) resulted in the loss of IL-18 processing in response to *H. pylori* stimulation (Fig. 4e, 4f). As the *H. pylori* T4SS is critical to induction of the NOD1 signalling pathway in AGS cells ^12, 14, 15^, we next studied IL-18 processing in response to isogenic *H. pylori* Δ*cag*PAI or Δ*cag*M strains, as well as to the mouse-adapted SS1 strain, all of which lack a functional T4SS ^12^. Lower levels of mature IL-18 production were observed in response to these strains when compared with the respective WT (251) and parental human clinical (10700) strain (Fig. 4e). Thus, *H. pylori* T4SS activation of the NOD1 signalling pathway is required for IL-18 processing in human gastric epithelial cells.

### NOD1 mediates IL-18 processing independently of RIPK2–NF-κB signalling

Classically, NOD1 activation results in the recruitment of the adaptor molecule RIPK2, induction of the NF-κB signalling pathway and upregulation of pro-inflammatory cytokine production ^12, 13, 14, 15^. We therefore investigated the role of RIPK2 in *H. pylori*-induced IL-18 processing in epithelial cells by small interfering RNA (siRNA) transfection or blocking its activity with the kinase inhibitor, WEHI-345 ^23^. siRNA-mediated KD of *RIPK2* was confirmed by measuring IL-8 production as a read-out for classical NOD1 pathway activation ^15^. Surprisingly, blocking RIPK2 signalling had no effect on IL-18 processing in response to *H. pylori* stimulation in AGS cells (Fig. 5a, 5b). In agreement with this finding, IL-18 processing was unaffected in both primary gastric epithelial cells and BMDMs from *Ripk2*^−/−^ mice, whereas epithelial cell production of the NOD1- regulated chemokines, Cxcl1/keratinocyte chemoattractant (KC) and Cxcl2/macrophage inflammatory protein 2 (MIP2) ^12, 24, 25, 26^, was reduced in response to *H. pylori* stimulation (Supplementary Fig. 6).

**Fig. 5.**
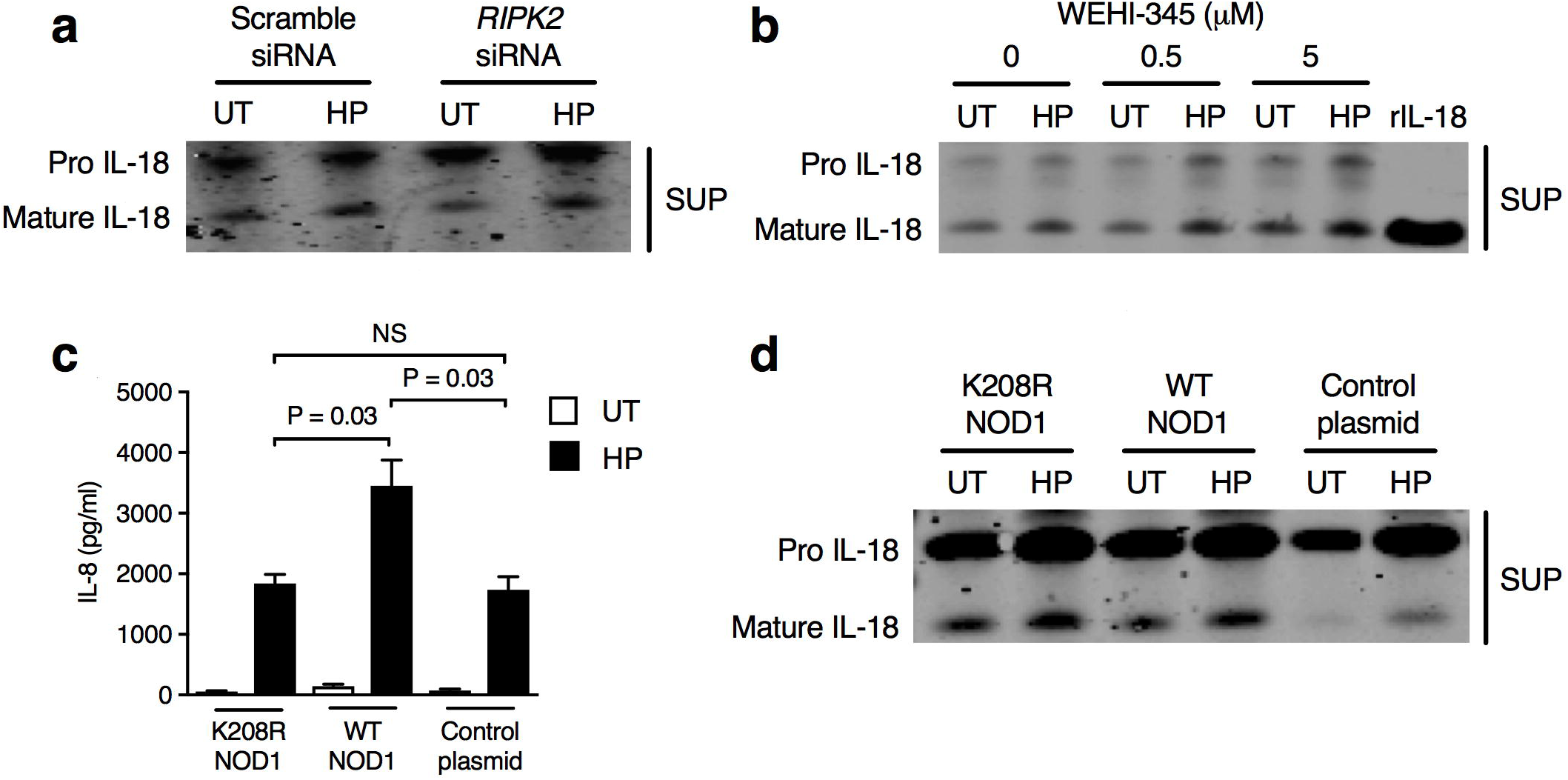
The NOD1 adapter protein RIPK2 is dispensable for NOD1-dependent Il-18 processing in epithelial cells. **a** Human AGS gastric epithelial cells were either pre-treated with scramble or RIPK2 siRNA then stimulated with *H. pylori* (HP) or left untreated (UT). IL-18 processing was assessed in culture supernatants (SUP). **b** AGS cells were pre-treated with varying concentrations of the RIPK2 inhibitor WEHI-345 then stimulated or not with *H. pylori*. **c, d** shNOD1 AGS cells were transfected with either a mutant form of NOD1 (K208R NOD1), WT NOD1 or the vector (control plasmid). IL-8 production (**c**) and IL-18 processing (**d**) was assessed in culture supernatants. Representative data for three independent experiments (**a-d**). Significance was determined by the unpaired two-tailed t-test (**c**). NS = not significant. Data correspond to the mean ± SEM.

To demonstrate that NF-κB activation by NOD1 is dispensable for IL-18 processing in epithelial cells, we restored NOD1 expression in sh*NOD1* AGS cells with plasmid constructs expressing either WT or a mutant form of NOD1 (K208R NOD1) that is unable to activate NF-κB signalling 19. As expected, NOD1 sh*NOD1* AGS cells transfected with plasmid encoding WT NOD1 exhibited significantly higher IL-8 responses to *H. pylori* stimulation than cells transfected with either the mutant NOD1 or empty control plasmids (Fig. 5c; *p* = 0.03). Conversely, transfection with either WT or mutant NOD1 plasmids was able to rescue IL-18 processing at comparable levels in sh*NOD1* AGS cells (Fig. 5d), suggesting that NOD1 mediates gastric epithelial IL-18 processing in an NF-κB independent manner. These findings suggest that *H. pylori* induces IL-18 processing via a non-classical type of NOD1 signalling pathway.

### NOD1 activates caspase-1 to promote IL-18 processing in epithelial cells

It is well established that caspase-1 is a key protease responsible for the cleavage of pro-IL-18 to its mature form in innate immune cells ^1, 16^. Therefore, to examine whether NOD1 mediates IL-18 processing in epithelial cells via caspase-1 activation, sh*NOD1* AGS cells that were transiently expressing yellow fluorescent protein (YFP)-labelled NOD1 were incubated with anti-caspase-1 antibody. As can be observed (Fig. 6a), NOD1 is normally proximal to the cell membrane under basal conditions, but moved towards the cytosol and formed yellow punctate stained aggregates with endogenous caspase-1, in response to *H. pylori* stimulation (Fig. 6a). This suggested that NOD1 may associate with caspase-1 upon activation by *H. pylori*. To demonstrate NOD1-mediated activation of caspase-1, we used the fluorescently labelled inhibitor of caspase (FLICA) reagent that binds irreversibly to active caspase-1. WT (sh*EGFP*) AGS cells exhibited enhanced staining with FLICA after stimulation with *H. pylori*, whereas staining was barely detected in sh*NOD1* AGS cells (Fig. 6b). Furthermore, we could only detect the mature, active form of caspase-1 (p20) in the culture supernatants of WT but not sh*NOD1* AGS cells in response to *H. pylori* stimulation (Fig. 6c). We also observed reduced levels of active caspase-1 in AGS cells stimulated with *H. pylori* mutant strains lacking either a functional T4SS (Δ*cag*PAI or Δ*cag*M) or defective in activating the NOD1 pathway (*slt^−^*; ^12^) (Fig. 6c). These results suggest that *H. pylori* factors required for NOD1 activation critically contribute to caspase-1 activation in gastric epithelial cells. Importantly, inactivation of caspase-1 with the inhibitor Z-YVAD resulted in a dose-dependent reduction of IL- 18 production in cells (Fig. 6d). Collectively, these data show that NOD1 sensing of *H. pylori* drives IL-18 processing via activation of caspase-1.

**Fig. 6.**
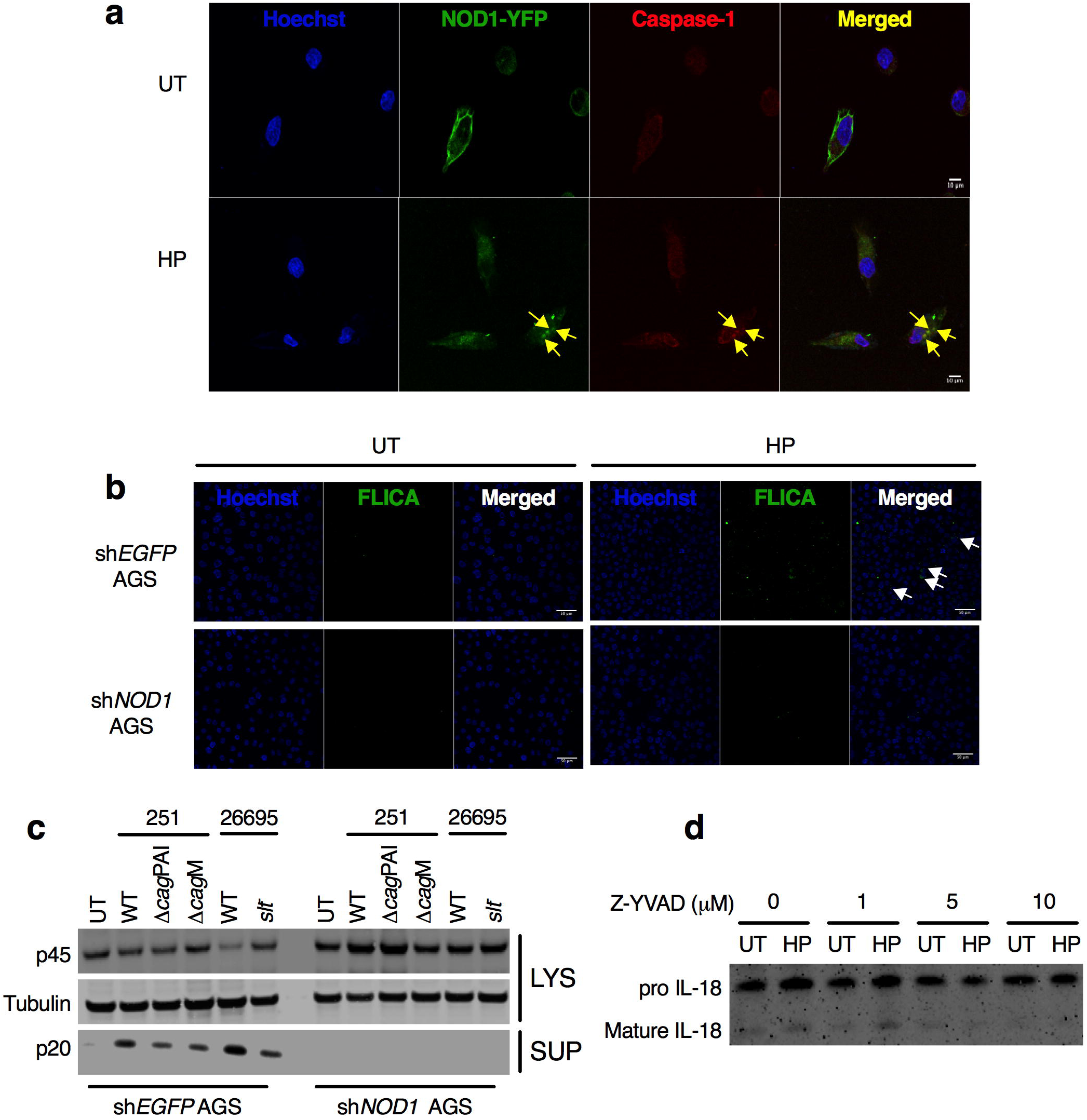
NOD1 activates caspase-1 to promote IL-18 processing in epithelial cells. **a** Human AGS gastric epithelial cells stably expressing shRNA to *NOD1* (sh*NOD1*) were transfected with YFP-labelled NOD1, stimulated with *H. pylori* (HP) or left untreated (UT), then incubated with an anti-caspase-1 antibody. In response to *H. pylori* stimulation, NOD1 and caspase-1 molecules can be observed co-localising (arrows). Cell nuclei were stained with Hoechst 33342, the images merged and the sections analysed by confocal microscopy. Scale = 10 µm. **b** AGS cells stably expressing shRNA to either *EGFP* (sh*EGFP*) or *NOD1* (sh*NOD1*) were stimulated with *H. pylori* or left untreated and then incubated with FLICA, a fluorescently labelled caspase inhibitor that binds irreversibly to caspase-1. Increased levels of fluorescence can be seen in the sh*EGFP* cells responding to *H. pylori* stimulation (arrows). Cell nuclei were stained with Hoechst 33342 and the images merged. Scale bar = 50 µm. **c** sh*EGFP* and sh*NOD1* AGS cells were left untreated (**UT**), or stimulated with the indicated *H. pylori* WT and mutant strains. Pro-(p45) and mature (p20) forms of caspase-1 were detected by Western blotting of cell lysates (LYS) and supernatants (SUP). **d** AGS cells were pre-treated with varying concentrations of the caspase-1 inhibitor Z-YVAD then stimulated or not with *H. pylori*. IL-18 processing was determined by Western blotting. Representative data for two independent experiments.

### NOD1 interacts with caspase-1 via CARD–CARD interactions independently of ASC

We next sought to elucidate potential NOD1-caspase-1 molecular interactions and their involvement in IL-18 processing. It is well established that NLRP3 and AIM2 need the adaptor protein, ASC, to recruit and activate caspase-1. However, as reported above (Supplementary Fig. 4), the AGS cell line does not produce detectable levels of ASC protein, suggesting that NOD1 may directly interact with caspase-1 via homotypic CARD–CARD interactions. This hypothesis was addressed using the fluorescence lifetime imaging microscopy-Förster resonance energy transfer (FLIM-FRET) technique in live cells. For this, we transfected sh*NOD1* AGS cells with mOrange-NOD1 plasmid (donor alone control) or both mOrange-NOD1 and the acceptor plasmid, encoding EGFP-caspase-1 (Fig. 7a). We found that *H. pylori* stimulation led to a significant reduction in the lifetime of the donor fluorochrome, indicative of quenching of the donor by the acceptor EGFP- caspase-1 due to close proximity (< 10 nm) of NOD1 and caspase-1 proteins (Fig. 7a; *p* < 0.0001). In contrast, the lifetime of the donor remained unchanged in unstimulated cells. These data confirm that NOD1 can directly interact with caspase-1 in response to *H. pylori* stimulation.

**Fig. 7.**
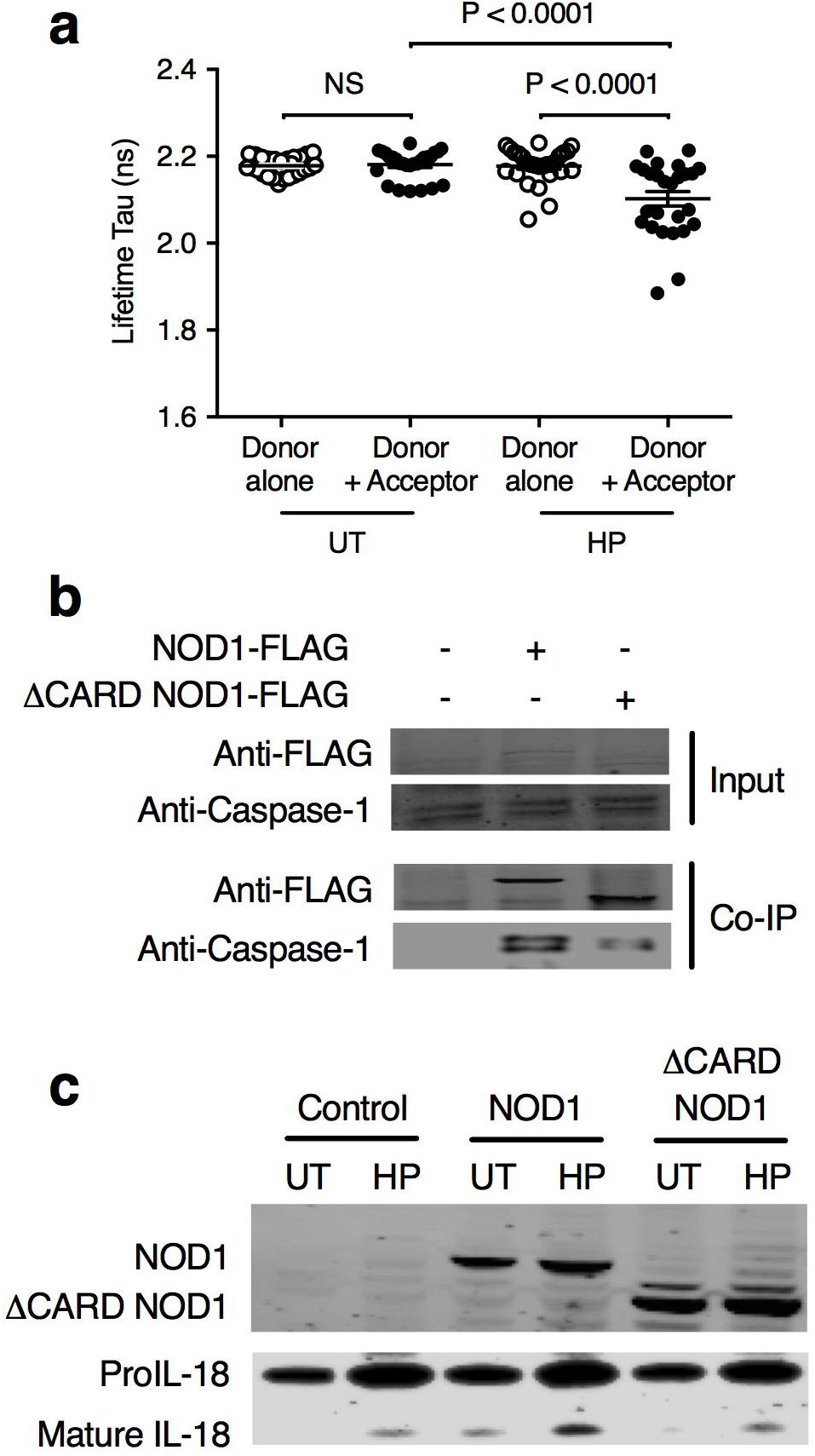
NOD1 interacts with caspase-1 via CARD–CARD interactions. **a** Human AGS gastric epithelial cells stably expressing shRNA to *NOD1* (sh*NOD1*) were transfected with mOrange-NOD1 plasmid (Donor alone) or both mOrange-NOD1 and EGFP-caspase-1 (Donor + Acceptor). Cells were then stimulated with *H. pylori* (HP) or left untreated (UT) and NOD1**–**caspase-1 interactions determined by the FLIM-FRET technique. A decrease in the lifetime of the donor fluorochrome, indicative of quenching of the donor by the acceptor, indicates close proximity (< 10 nm) between NOD1 and caspase-1 proteins. **b** *NOD1* KO AGS cells, generated by CRISPR/Cas9 gene editing, were transfected with FLAG-tagged plasmids encoding either full-length NOD1 (NOD1-FLAG) or NOD1 lacking a CARD domain (ΔCARD NOD1-FLAG) ^59^. Proteins in input and immunoprecipitated samples (Co-IP) were detected using anti-FLAG and -caspase-1 antibodies. **c** *NOD1* KO AGS cells were transfected with either full-length NOD1 (NOD1) or CARD-deficient NOD1 (ΔCARD NOD1), then stimulated with *H. pylori* or left untreated. Proteins were detected in cell lysates using anti-NOD1 or anti-IL-18 antibodies on cell lysates or supernatants, respectively. Representative data for two independent experiments. Data were pooled from two independent experiments and significance was determined by the unpaired t- test (**a**). Data correspond to the mean ± SEM.

To further examine whether NOD1 interacts with caspase-1 via homotypic CARD–CARD interactions, we performed co-immunoprecipitation assays in which *NOD1* KO AGS cells with FLAG-tagged plasmids encoding either full-length NOD1 or NOD1 lacking a CARD domain (ΔCARD NOD1). Full-length and truncated NOD1 proteins were pulled down using anti-FLAG antibodies. Analysis of the resulting immune-complexes revealed that full length NOD1, but not NOD1 lacking a CARD domain, was able to strongly associate with endogenous caspase-1 (Fig. 7b). Moreover, reconstitution of NOD1 expression with full-length but not CARD-deficient NOD1 rescued *H. pylori*-induced IL-18 processing in these cells (Fig. 7c), thus highlighting the important role of the CARD in mediating IL-18 processing. Collectively, the results demonstrate that NOD1 sensing of *H. pylori* in epithelial cells activates caspase-1 via direct CARD–CARD interactions and independently of ASC, resulting in the processing of IL-18.

### NOD1 maintains epithelial homeostasis in response to *H. pylori* infection

As IL-18 appeared to be important for protecting tissues against excessive pathology during chronic *H. pylori* infection *in vivo* (Fig. 2, Supplementary Fig. 1), we next asked whether NOD1-mediated IL-18 secretion may play a role in regulating gastric epithelial cell survival responses. For this, we performed *in vitro* assays including MTT assay and Annexin V/propidium iodide (PI) staining to assess cell proliferation and apoptosis, respectively. sh*NOD1* KD AGS cells exhibited significantly higher levels of both cell growth and apoptosis in response to *H. pylori* stimulation, when compared with sh*EGFP* AGS cells (Fig. 8a, 8b; *p* = 0.001 and *p* = 0.01, respectively). Differences in apoptosis were not observed in cells stimulated with the apoptotic-inducing agent, etoposide (Fig. 8b). These data suggest that NOD1 is involved in regulating both epithelial cell growth and death in response to *H. pylori* stimulation. To confirm these findings, we established a 3-D gastric organoid model from *Nod1^+/+^* and *Nod1^−/−^* mice. *H. pylori* bacteria were added to the lumen of the organoids by microinjection (Fig. 8c). Consistent with data for AGS cells (Fig. 2a, b), *Nod1*^−/−^ organoids exhibit higher levels of cell proliferation and apoptosis, as compared with *Nod1*^+/+^ organoids (Fig. 8d, 8e; *p* = 0.02 and *p* = 0.008, respectively). Collectively, these results show that NOD1 is important for maintaining homeostasis in gastric epithelial cell turnover in response to *H. pylori* infection.

**Fig. 8.**
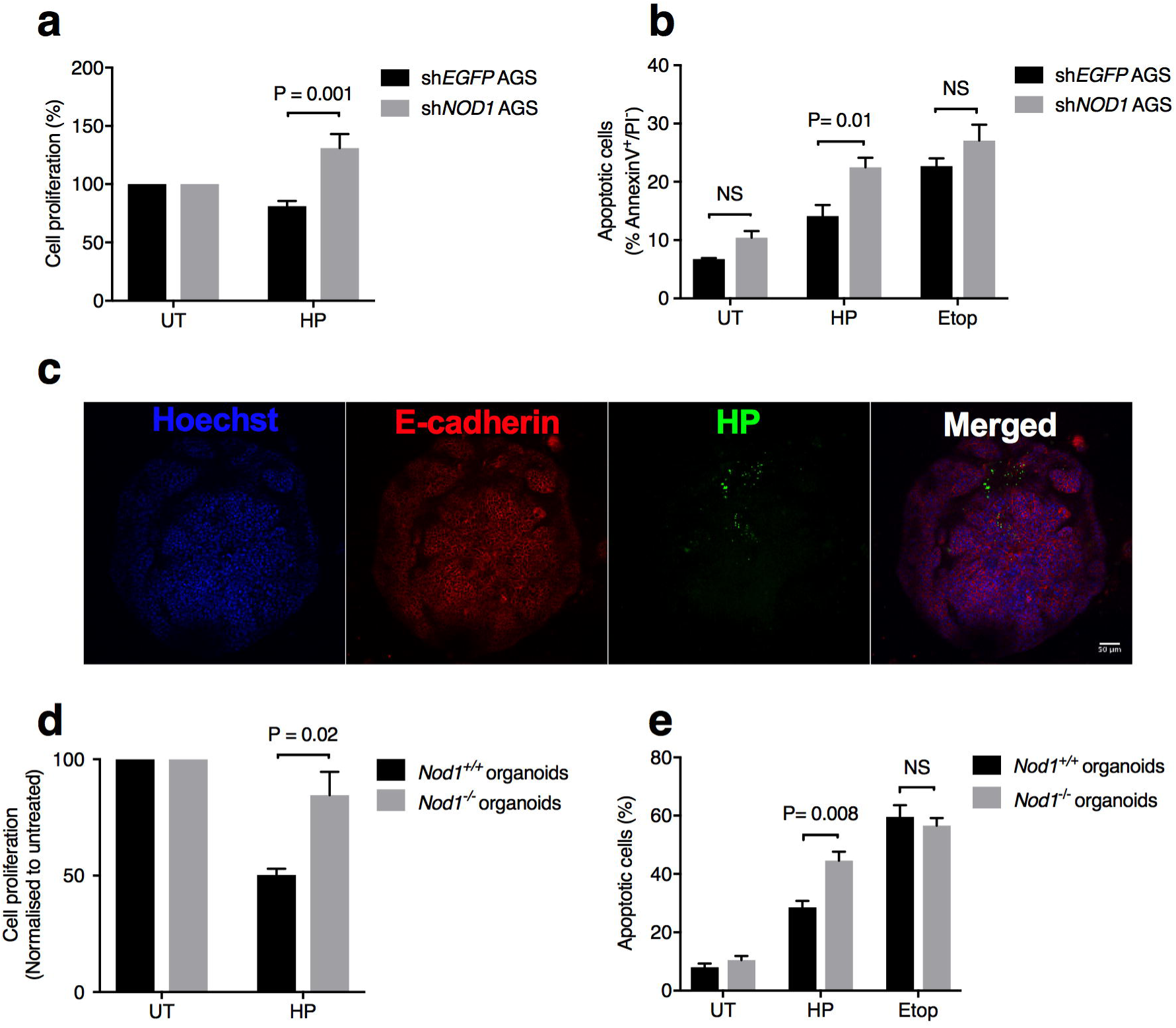
NOD1 maintains epithelial homeostasis in response to *H. pylori* infection. **a**, **b** Human AGS gastric epithelial cells stably expressing shRNA to either *EGFP* (sh*EGFP*) or *NOD1* (sh*NOD1*) were stimulated with *H. pylori* (HP) or left untreated (UT). Cell proliferation (**a**) and apoptosis (**b**) were determined using the MTT assay and Annexin V/PI staining, respectively. As a control, cells were treated with the apoptotic-inducing agent, etoposide (5 μM). **c-e** Gastric organoids were isolated from *Nod1^+/+^* and *Nod1^−/−^* mice, then either left untreated (UT) or treated with *H. pylori* (HP) or etoposide. *H. pylori* bacteria (green) were microinjected into the lumen of organoids consisting of E-cadherin^+^ cells (red) (**c**). Cell nuclei were stained with Hoechst 33342, the images merged and the sections analysed by confocal microscopy. d, e Gastric organoid cultures from *Nod1^+/+^* and *Nod1^−/−^* mice were assessed for changes in cell proliferation (**d**) and apoptosis (**e**), using Presto Blue assay and Annexin V/propidium iodide (PI) staining, respectively. Representative data for two independent experiments (**a-e**). Significance was determined by the unpaired two-tailed t- test. Data correspond to the mean ± SEM.

## Discussion

Current understanding of canonical inflammasome functions has largely been shaped by studies in haematopoietic cells and in particular, the processing of the pro-inflammatory cytokine, IL-1β. In contrast, the functions of inflammasomes in non-haematopoietic cell lineages, including epithelial cells, are much less well understood ^3^. The inflammasome molecules AIM2, NLRP3, NLRP6 and NLR Family Apoptosis Inhibitory Protein (NAIP)/NLRC4 have so far been reported to play important roles in epithelial cells by protecting against infection and maintaining tissue homeostasis ^3, 27, 28, 29, 30, 31^. An important distinction between epithelial inflammasomes and those in mononuclear phagocytes is that the former are primarily involved in IL-18 production, whereas the latter are a major source of IL-1β ^3^. IL-18 was initially identified as promoting intestinal inflammation, but there is now overwhelming evidence to support the contention that this cytokine can also play protective roles against colitis ^3, 29, 32, 33, 34^. In the current study, we show that IL-18 plays a protective role against the development of pre-neoplastic changes in the stomach due to chronic *H. pylori* infection. Moreover, we identify a new type of inflammasome as being important for epithelial cell production of mature, biologically active IL- 18.

Intestinal epithelial cells constitutively produce IL-18, suggesting that epithelial inflammasomes have a “ready-to-go” phenotype that allows rapid responses to pathogens and other damage to the gut ^3, 29^. In agreement with this suggestion, we (Fig. 4, 5) and others ^4, 5, 6^ have established that IL- 18 is expressed constitutively in the gastric mucosa, with gene expression and protein levels upregulated in response to *H. pylori* infection. Nevertheless, the role of IL-18 responses in *H. pylori* infection and associated diseases is unclear. A gene polymorphism study identified certain haplotypes as possibly being associated with susceptibility to *H. pylori* infection in the Korean population ^35^, but functional data for IL-18 responses were not provided. Other studies could not find any correlations between IL-18 haplotype, phenotype and clinical outcome, including gastric cancer ^4, 36^. Conversely, a correlation was found between certain IL-18 haplotypes, higher gastric IL-18 levels and more severe mononuclear cell infiltration in *H. pylori*-infected subjects ^4^. Nevertheless, this does not prove causality and may simply reflect IL-18 production by the mononuclear cells recruited to the site of infection. Yamauchi *et al.* isolated mononuclear and epithelial cells from the gastric mucosa and showed that both cell types produce IL-18 in response to *H. pylori* stimulation *ex vivo* ^6^, but was unable to determine the relative contributions of these cells to IL-18 production *in vivo*. Our data provide unequivocal evidence that epithelial cells are a major source of IL-18 production in the stomach during *H. pylori* infection (Fig. 1c, d; 2, a, b). Using primary and immortalised epithelial cells, we show that these cells can also process IL-18 to its mature form (Fig. 4b, d-f). It has been suggested that studying the balance between pro- and mature IL-18 forms may be more informative in regards to understanding the role of this cytokine in inflammatory diseases ^5^. Further studies are therefore warranted to investigate both the localisation and form of IL-18 present in the gastric tissues from *H. pylori*-infected subjects presenting with diseases of varying severity.

It has been reported by several groups that *H. pylori* induces NLRP3 inflammasome activation and IL-1β responses in dendritic cells, neutrophils and macrophages/monocytes ^37, 38, 39, 40, 41, 42^. *H. pylori* was shown to upregulate IL-18 production in these immune cells ^7, 38, 39, 40^ via an NLRP3- dependent mechanism ^39, 40^. Similarly, we observed NLRP3-dependent regulation of IL-1β and IL-18 responses in BMDMs (Supplementary Fig. 2), however, no role could be found for this inflammasome, nor for its adaptor ASC, in IL-18 production by primary mouse gastric epithelial cells (Fig. 3a). *H. pylori* upregulated IL-18 production and processing in the human AGS gastric epithelial line (Fig. 4e, f, 5d) which, contrary to previous reports ^43, 44^, does not appear to express either NLRP3 or ASC (Supplementary Fig. 4 and unpublished data). Another study also reported that AGS cells do not produce ASC ^6^. Consistent with these data, no differences in *H. pylori* bacterial loads or resulting inflammation were observed in *Nlrp3*^−/−^ or *Pycard*^−/−^ mice when compared with WT animals (Fig. 3b-d). One group reported increased bacterial loads and decreased gastric inflammation in *Nlrp3*^−/−^ mice ^42^, whereas the complete opposite was reported in another study ^38^. The reasons for the divergent results between these three studies are unclear but may stem from differences in the composition of the mouse microbiota and not to the bacterial strains used, as the latter two studies used the same strain *i. e. H. pylori* 10700 (PMSS1). Notwithstanding the different findings, the data presented here clearly establish that the canonical NLRP3/ASC inflammasome is not required for gastric epithelial cell production of IL-18 responses to *H. pylori* infection (Fig. 3; 4e, f). In addition, the inflammasome protein NLRC4 does not seem to play a significant role in the infection (Fig. Supplementary Fig. 3g, h). This finding is compatible with the known inability of *H. pylori* ^42^ or its flagellin ^45^ to promote caspase-1 processing and IL-1β secretion in macrophages.

A key and novel discovery of the current work is the observation that NOD1 interacts with caspase-1 to mediate *H. pylori* induction of IL-18 processing in epithelial cells. Interestingly, this IL-18 processing occurs independently of CARD-CARD interactions between NOD1 and its adapter molecule, RIPK2 (Fig. 5). Early studies in the field reported NOD1 interactions with caspase-1 ^19, 46^, resulting in IL-1β secretion in cells ^46^. A caveat to those studies, however, is that they were undertaken using over-expression approaches which can be susceptible to experimental artefacts. We now provide several lines of supportive evidence to show that NOD1 interacts with caspase-1 to mediate tissue protective IL-18 responses in epithelial cells. Firstly, we have confirmed the findings of previous investigations by performing similar over-expression and classical protein “pulldown” experiments (Fig. 7b, c) and used fluorescence and FLIM-FRET imaging to show close interactions between NOD1 and caspase-1 in response to *H. pylori* stimulation (Fig. 6a, b; Fig. 7a). Secondly, we demonstrate that caspase-1 processing is itself dependent on NOD1 activation (Fig. 4, 6c). Thirdly, we show that IL-18 production is abrogated in primary gastric epithelial cells from *Casp1*^−/−^ mice (Fig. 3a) and in cells pre-treated with the caspase-1 inhibitor, Z-YVAD (Fig. 6d). Finally, *Casp1*^−/−^ mice infected with *H. pylori* reproduce aspects of the severe pathology observed in *Il18*^−/−^ animals (Fig. 3b-d) and is consistent with the enhanced inflammatory changes also observed in *Casp1*^−/−^ mice infected with *H. felis* ^7^.

NLRs were originally identified for their roles in host defence and inflammation. However, more recent work in colitis models has shown that these proteins also play important roles in tissue repair and the maintenance of homeostasis. NLRs were shown to mediate these protective effects via various mechanisms, including the production of antimicrobial peptides, control of the gut microbiota ^28, 30^, prevention of tumour development ^27, 47^ and regulation of NF-κB and MAPK signalling ^48^. A key downstream mediator of these tissue protective responses is IL-18 ^28, 30, 49^. Indeed, administration of IL-18 to *Casp1*^−/−^ mice protected animals from tissue damage and death caused by dextran sulphate sodium treatment ^49^. We propose that IL-18 plays a similar protective role against tissue damage caused by chronic *H. pylori* infection. Our data show that NOD1 is an important regulator of bioactive IL-18 production by epithelial cells in response to the infection and, moreover, that this NLR plays a protective role in tissue homeostasis through its control of cell proliferation and apoptosis (Fig. 8). Interestingly, it was shown in the Mongolian gerbil model that treatment with a NOD1 agonist prior to *H. pylori* challenge could suppress the development of inflammation and carcinogenesis in animals ^26^.

In conclusion, we have identified a new NOD1 inflammasome pathway that mediates different cellular responses to those regulated by classical NOD1→RIPK2→NF-κB signalling. Similar to other NLR inflammasomes, activation of NOD1 signalling via a single stimulus (*i. e. H. pylori*) promotes both inflammatory and tissue repair responses in cells. These responses occur simultaneously in cells *in vitro*, but may be temporally regulated *in vivo*, particularly over long periods, such as chronic *Helicobacter* infection. This regulation may occur via the actions of CARD-containing proteins which have been shown to differentially regulate inflammatory and cell death pathways ^50^. Indeed, it was shown that the CARD-containing protein, CARD6, could selectively modulate NOD1/NF-κB signalling, independently of caspase-1/IL-1β secretion ^51^. Future studies are therefore required to determine whether NOD1 can be selectively activated to produce tissue protective IL-18 responses that are beneficial to the host. It also remains to be determined whether other Gram-negative pathogens that colonise mucosal surfaces are able to activate NOD1 signalling to promote tissue repair and possibly, even anti-tumourigenic responses.

## Methods

### Mouse strains, cell lines and bacterial strains

*Casp1^−/−^(Casp1^−/−^Casp11^−/−^*) ^52^, *Il18^−/−^* ^52^, *Nlrp1^−/−^*^53^*, Nlrp3^−/−^* ^54^, *Nlrc4*^−/− 55^, *Pycard*^−/− 52^*, Ripk2*^−/− 23^ and WT C57BL/6J mice were bred as litter mates at the Walter Eliza Hall Institute of Medical Research (WEHI). *Nod1^−/−^* ^15^, *Nod1*^fl/fl^ and *Nod1*^fl/fl^ x LysM-Cre mice (Supplementary Fig. 7) and WT mice were bred at the Animal Research Facility, Monash Medical Centre (MMC). KO mice had been backcrossed at least ten generations on the C57BL/6J background and maintained under specific pathogen-free conditions in micro-isolator cages with access to food and water *ad libitum*. Experiments were performed using age- and sex-matched animals. Mouse primary gastric epithelial cells ^15^ and BMDMs ^52^ were isolated, as described previously. All animal procedures complied with guidelines approved by the Animal Ethics Committees at the WEHI (2014.004, 2011.014, 2008.022) and MMC (MMCA/2015/43), respectively.

The human AGS gastric cancer cells stably expressing shRNA to either *EGFP* (sh*EGFP*) or *NOD1* (sh*NOD1*) were generated as described previously ^22^. Human *NOD1* KO and control AGS cells were generated by CRISPR Cas/9 technology and their KO status validated previously ^15^. AGS cells were maintained in complete RPMI medium supplemented with 10% (v/v) foetal calf serum (FCS), 1% (w/v) L-glutamine and 1 % (w/v) penicillin/streptomycin (Gibco; ThermoFisher Scientific, VIC, Australia).

The mouse GSM06 GEC line (Riken Cell Bank RCB1779) was grown as described previously 20. Briefly, the cells were maintained in Dulbecco’s modified Eagle medium–nutrient mixture/F-12 medium, supplemented with 10% FCS, 1% (w/v) insulin/transferring/selenite (Gibco) and 10 ng/ml epidermal growth factor. Cells were grown in 5% CO_2_ at the permissive temperature of 33°C, then moved to 37°C prior to experiments.

*H. pylori* strains 251 (WT, Δ*cag*PAI, Δ*cagM*; ^12, 14^), 26695 (WT, *slt^−^*;^12^), 245m3 ^56^, B128 7.13 ^57^ the mouse-adapted SS1 strain and its clinical progenitor isolate 10700 (also known as PMSS1) ^58^ were routinely grown on blood agar medium containing selective antibiotics at 37°C under microaerobic conditions ^20^.

### Mouse *H. pylori* infection

Mice (6-8 week-old) were inoculated via oral gavage with 10^8^ colony-forming units of *H. pylori* strain SS1 ^20^, B128 7.13 ^57^ or 245m3 ^20^. Bacterial viability and numbers were determined by agar plate dilution. Stomachs and spleens were harvested at 1 and 8 weeks p.i. Gastric tissues were subjected to homogenisation using a gentleMACS™ Dissociator (Miltenyi Biotec, NSW, Australia). Bacterial infection status was confirmed by bacteriological culture ^20^. Gastric homogenates were centrifuged at 13,000 × g at 4°C and the supernatants collected for analysis by the Qubit™ protein assay (ThermoFisher Scientific) and ELISA.

BM reconstitution experiments were performed by subjecting recipient mice to two 5.5-Gy doses of irradiation given 3 hr apart and then injecting 1 × 10^6^ donor bone marrow cells via intravenous injection ^52^. At 6 weeks post-transfer, recipient mice were infected with *H. pylori* SS1 and sacrificed at 5 and 18 weeks p.i.

### Inhibitor treatment

Cells were pre-treated with ML130 (5 μM; CID-1088438; Tocris, UK) ^21^, WEHI-345 (0, 5 or 10 μM) ^23^ or Z-YVAD-fmk (0, 1, 5 and 10 μM; 21874; Merck, Australia) for 1 hr prior to stimulation with *H. pylori*, then maintained in inhibitor-containing medium throughout the experiment.

### Histological analysis

Gastric tissues were formalin-fixed and embedded in paraffin. Tissues were stained with Mayer’s Haematoxylin & Eosin or Periodic acid-Schiff (PAS)-Alcian Blue (AB) stains and imaged at 20 x magnification using a bright-field microscope (Nikon). Inflammatory scores for neutrophils and lymphocytes were graded in a blinded fashion (JSP) according to a previously described grading scheme ^20^. The mucosal thickness of stomach tissues was determined in a blinded fashion (JE) on sections that had been digitised using an Aperio slide scanner (Leica Biosystems, VIC, Australia) and ImageScope software (Leica Biosystems). Three measurements were taken for each tissue section, with 3-4 sections available per stomach sample.

#### Immunofluorescence

Antigen retrieval was performed by immersion in 10 mM sodium citrate buffer (pH 6) and boiling for 8 min in a microwave. Sections were washed twice with PBS and immersed in blocking buffer (5% horse serum, 1% Triton X100). After 1 hr, sections were incubated overnight at 4°C with an antibody mix containing rat anti-mouse IL-18 (5 μg/ml; D047-3; R&D Systems, MN, USA) and rabbit anti-EpCAM (Clone ab71916, Abcam). An isotype control for IL-18 staining was performed using mouse IgG (Dako). Samples were washed thrice with blocking buffer and incubated for 2 hr with anti-rat Alexa Fluor® 488- and anti-rabbit Alexa Fluor® 594-conjugated secondary antibodies (ThermoFisher Scientific, 1:400). After washing, cell nuclei were stained with Hoechst 33342 (ThermoFisher Scientific; 1:1000) for 5 min before mounting. Imaging was performed on a confocal microscope (Nikon Instruments Inc., NY, USA). All images were captured using the same settings.

#### Cytokine ELISA

Mouse IL-1β (BD Biosciences, NSW, Australia), IL-18 (R&D Systems), Cxcl1/KC (R&D Systems), Cxcl2/MIP2 (R&D Systems), human IL-18 (R&D Systems) and CXCL8 (BD Biosciences) were quantified by ELISA, as per the manufacturers’ instructions.

#### Cell sorting

Single-cell suspensions were prepared from stomach tissues of *H. pylori* infected mice. Briefly, whole stomachs were finely minced in disassociation buffer (2% FCS in Hank’s Balanced Salt Solution without calcium and magnesium (Life Technologies) and 5 mM EDTA) and incubated at 37°C with shaking at 180 rpm for 30 min. Tissues were passed through 70 μm cell strainers (Falcon, ThermoFisher Scientific) and the filtrates were collected. The remaining pieces of tissues were further digested at 37°C with shaking at 180 rpm for 45 min in 2% FCS in RPMI (Life Technologies), containing 1 mg/ml collagenase Type 1 (Life Technologies), 0.4 units Dispase (Life Technologies), and 0.01 mg/ml DNase (Roche). Samples were passed through cell strainers (70 μm) and the filtrates combined and centrifuged at 630 × g for 10 min at RT. Cell pellets were resuspended in FACS buffer (5% FCS in PBS, Life Technologies). After adding Fcγ receptor block (1:100 dilution; BD Biosciences) and incubating on ice for 10 min, dead cells were stained with PI (0.5 μg/ml; Life Technologies) and incubated for 30 min at 4°C with antibodies specific for the following markers: CD45.2-PE-Cy7 (1:200 dilution; BD Biosciences); EpCAM-APC (1:100 dilution; eBioscience, CA, USA). Viable CD45.2^+^ EpCAM^−^ and CD45.2^−^EpCAM^+^ cells were sorted (FACS Aria Fusion, Beckman Coulter), verified for cell purity and the RNA isolated using Trizol (Life Technologies).

### Cell co-culture assay

*H. pylori* bacteria were grown in BHI broth (Oxoid, ThermoFisher Scientific) supplemented with 10% (v/v) heat-inactivated foetal calf serum (ThermoFisher Scientific) and antibiotics, in a shaking incubator for 16-18 hr at 37°C under microaerobic conditions ^20^. Bacteria were pelleted and washed twice with phosphate-buffered saline (PBS) by centrifugation at 4,000 × g for 10 min at 4°C prior to resuspension in RPMI medium for co-culture assays. Viable counts were performed by serial dilution of bacterial suspensions on blood agar plates.

Cells were seeded in 12-well plates at 1 × 10^5^ cells/ml and allowed to grow overnight. The culture medium was removed and replaced with serum- and antibiotic-free RPMI medium prior to stimulation with bacteria. Cells were incubated with *H. pylori* WT or isogenic mutant strains at a multiplicity of infection (MOI) of 10:1. The bacteria were removed after 1 hr. Culture supernatants and lysates were collected at 24 hr post-stimulation for ELISA and Western blot analyses. FLICA staining was performed as per the manufacturer’s instructions (ImmunoChemistry Technologies, MN, USA). Cell proliferation was determined using the MTT assay (Promega) on adherent AGS cells, while apoptosis was determined in AGS cells detached from culture plates by the addition of 0.25% Trypsin and 1 mM EDTA for 2 min. The isolated cells were subjected to Annexin V/PI (Life Technologies) staining and analysed by Flow cytometry.

#### Cell transfection

DNA plasmids (Caspase-1-GFP and NOD1-mOrange) ^59^ or specific silencer and negative control siRNA (40 μmol of each; ThermoFisher Scientific) were diluted in Opti-MEM medium (Gibco) supplemented with 2 μl Lipofectamine® 2000 reagent (ThermoFisher Scientific). The mixtures were incubated at room temperature for 20 min then added drop-wise to each well of 12-well plates containing 10^5^ AGS cells. Transfected cells were incubated at 37°C for 24 hr prior to co-culture with bacteria.

#### FLIM-FRET analysis

AGS cells were co-transfected with caspase-1 GFP (donor) and NOD1-mOrange (acceptor) before either *H. pylori* stimulation or control. FLIM data was recorded on an Olympus FV1000 confocal microscope equipped with a PicoHarp300 FLIM extension and 485 nm pulsed diode (PicoQuant, Germany). The pixel integration time for FLIM imaging was maintained at 40 µs/pixel and fluorescence lifetime histograms were accumulated to at least 20,000 counts to ensure sufficient data collection for FLIM-FRET analysis. Data was collected with photon count rates at 5% of the laser repetition rate to prevent pileup. FLIM-FRET analysis was performed using Symphotime 64 software (PicoQuant, Germany), with individual cells selected as regions of interest (ROIs) for analysis. In each experiment, ten random fields of view were recorded for each treatment and a minimum of five cells from each field were analysed. The fluorescence lifetime decay of each cell was deconvolved using the measured instrument response function (IRF), then fitted with a bioexponential decay. The amplitude-weighted average lifetime was obtained from each fit and averaged over all cells for each condition.

#### Co-immunoprecipitation

Cells were seeded at 2 × 10^5^ cell/ml in 10 cm dishes and left to rest overnight. DNA (4 μg) of plasmid constructs encoding either FLAG-tagged full-length NOD1 (NOD1-FLAG) or NOD1 lacking a CARD domain (ΔCARD NOD1-FLAG) ^59^ was diluted in 1 ml of Opti-MEM medium (Gibco) supplemented with 15 μl Lipofectamine® 2000 reagent (ThermoFisher Scientific). The mixtures were incubated at room temperature for 20 min, then added to each dish. After stimulation with bacteria, cells were harvested and lysed in low-stringency lysis buffer (150 mM NaCl, no detergent). Lysate supernatants were pre-cleared by adding 10 µl of protein A/G agarose beads (Santa Cruz) and rotating samples at 4°C for 30 min. After collecting supernatants, 25 μl of agarose beads and anti-FLAG antibody (1 μg; F1804; Sigma Aldrich, MO, USA) were added. The mixtures were rolled at 4°C for 2-3 hr and washed with low stringency buffer. Beads were collected and resuspended in LDS sample buffer (50 μl; Life Technologies) and boiled at 95.4°C for 5 min, prior to Western blot analysis.

#### Western blot analyses

Cell culture supernatants were harvested and concentrated with StrataClean resin beads (Agilent Technologies, CA, USA), as per the manufacturer’s instructions. Cell lysates were prepared using NP-40 lysis buffer (ThermoFisher Scientific), supplemented with complete protease and phosphatase inhibitors (Roche). Total protein lysates (50 µg) were resuspended in Laemmli buffer (30 μl; ThermoFisher Scientific). All samples were heated at 98°C for 10 min, loaded onto NuPAGE® 4-12% gels and run at 120 V in 1 X MES buffer (ThermoFisher Scientific). The separated proteins were transferred onto membranes using the iBlot® transfer system (ThermoFisher Scientific), as per the manufacturer’s instructions. The membranes were blocked using Odyssey® blocking buffer (Odyssey; LI-COR, NE, USA). Membranes were incubated overnight at 4°C, with rat anti-mouse IL-18 (1 μg/ml; D047-3; R&D Systems), rabbit anti-human IL-18 (0.2 μg/ml; sc-7954; Santa Cruz Biotechnology), mouse anti-human caspase-1 (0.1 μg/ml; sc56036; Santa Cruz Biotechnology), rabbit anti-Asc (1 μg/ml; clone AL177; AdipoGen, CA, USA), rabbit anti-human NLRP3 (1 μg/ml; Clone D2P5E; Cell Signaling, CA, USA) or mouse anti-FLAG (1 μg/ml; F1804; Sigma). Membranes were washed in PBS-0.05% (v/v) Tween 20 and incubated for 2 hr with goat-anti-mouse secondary antibody-Alexa Fluor^®^ 680 conjugate or goat anti-rabbit secondary antibody-Alex Fluor^®^ 800 (0.67 ng/ml; ThermoFisher Scientific). Membranes were washed and developed on the Odyssey Infrared Imaging System (LI-COR). As loading controls, lysate samples were probed with rat-anti-human Tubulin (0.9 ng/ml; Rockland, PA, USA), followed by goat-anti-rat secondary antibody-Alexa Fluor^®^ 800 conjugate (0.33 ng/ml; ThermoFisher Scientific).

#### Quantitative PCR (qPCR) analyses

RNA was extracted from cells using the PureLink® RNA mini kit (ThermoFisher Scientific). cDNA was generated from 500 μg of RNA using the Tetro cDNA synthesis kit (Bioline, NSW, Australia), as per the manufacturer’s instructions. cDNA (1 µl) was used per reaction, consisting of 5 X Green GoTaq Flexi Buffer (Promega), dNTP mix (0.2 mM per dNTP) and primer (1 μM). Oligonucleotide sequences were as follows: *RNA18S1*, Fwd-CGGCTACCACATCCAAGGAA and Rev-GCTGGAATTACCGCGGCT; *PYCARD*, Fwd-TCACCGCTAACGTGCTG and Rev-TGGTCTATAAAGTGCAGGCC; *NLRP3*, Fwd-GTGTTTCGAATCCCACTGTG and Rev-TCTGCTTCTCACGTACTTTCTG; *Rn18s*, Fwd-GTAACCCGTTGAACCCCATT and Rev-CCATCCAATCGGTAGTAGCG; *Cxcl2*, Fwd-AACATCCAGAGCTTGAGTGTGA and Rev-TTCAGGGTCAAGGCAAACTT; *Il1b*, Fwd-ACGGACCCCAAAAGATGAAG and Rev-TTCTCCACAGCCACAATGAG; *Il18*, Fwd-GCCTCAAACCTTCCAAATCAC and Rev-GTTGTCTGATTCCAGGTCTCC. Denaturation was performed at 95°C for 15 seconds, then 20 successive cycles of amplification (95°C, 15 sec; 60°C, 1 min) and extension at 72°C for 30 sec. PCR products were run at 100 V for 20 min on 2% (w/v) agarose gels and stained with SYBR Safe DNA Gel Stain (Invitrogen; Carlsbad, California, USA). Gels were visualised in a UV trans-illuminator and imaged using Quantum-Capture software.

For qPCR, reactions consisted of synthesised cDNA (4 μl, 1:10 dilution), SYBR® Green qPCR MasterMix (5 μl; ThermoFisher Scientific) and primer (1 µl, 1 μM). Assays were performed in an Applied Biosystems™ 7900 Fast Real-Time PCR machine (ThermoFisher Scientific), using the following program: 50°C, 2 min, followed by 95°C, 10 min, then 40 successive cycles of amplification (95°C, 15 sec; 60°C, 1 min). Gene expression levels were determined by the Delta-Delta Ct method using the *18S rRNA* gene as a reference.

#### Organoid cultures and microinjection

Gastric organoids were generated using an adaptation of previously described methods ^60^. Briefly, mouse stomachs were dissected, cut open along the greater curvature, and washed thoroughly with PBS. The outer muscle layer was removed using fine-tipped forceps and the forestomach discarded. The remaining tissue was cut into 2 mm pieces and washed in cold PBS before digestion with 10 mM EDTA for 4 hr at 4°C. To isolate glands, tissue fragments were vigorously suspended in cold PBS using 10 ml pipettes and the supernatants were collected. This procedure was repeated a total of three times. Crypts were pelleted by centrifugation at 240 × g for 5 min, at 4°C. Pellets were resuspended in PBS, passed through cell strainers (70 µm; Falcon) and centrifuged. Supernatants were discarded and the pellets containing the glands were resuspended in Matrigel (Corning). Matrigel was seeded into each well of a 24-well plate (Nunc) and incubated for 10 min at 37°C until solidified. Crypt culture media (DMEM/F12 (Gibco), B27 (Gibco), Glutamax (Gibco), N2 (Gibco), 10 mM HEPES (Gibco), N-acetyl cysteine (1.25 µM; Sigma Aldrich), EGF (50 ng/ml; Peprotech), Noggin (100 ng/ml; Peprotech), Gastrin (10 nM; Sigma Aldrich), FGF10 (100 ng/ml; Peprotech), 10% (v/v) R-spondin 1-conditioned media, and 50% (v/v) Wnt3a-conditioned media) was added to each well. Established organoids were microinjected with 20 nl of overnight culture *H. pylori* suspension (10^7^ bacteria/ml) using the IM-300 pneumatic microinjector (Narishige). The presence of bacteria in the organoids was confirmed by immunofluorescence staining with rabbit anti-*H. pylori* antibody (in-house) and anti-cadherin (5 μg/ml; clone DECMA-1, Abcam), followed by the appropriate secondary antibodies. Organoid viability was assessed using PrestoBlue reagent (Life Technologies). Cell proliferation and apoptosis in the organoids were assessed using the MTT assay and Annexin V/PI straining, respectively.

## Acknowledgements

The authors thank: L. Inglis and L. Scott (WEHI) for animal husbandry; W. Carter (WEHI) for assistance with biosafety compliance; Assoc Prof. S. Bedoui (The University of Melbourne) for supplying the *Nlrc4*^−/−^ mice; and Mr Dirk Truman (Monash Gene Targeting Facility, Monash University) for designing the Nod1 targeting strategy and generating the heterozygote animals. The authors acknowledge assistance from the Monash BDI Organoid Program, Monash Biomedicine Discovery Institute, Victoria, Australia. Drs F. Cribbin and R. Smith (The Hudson Institute) are thanked for editorial assistance. The research in RLF’s laboratory was supported by project grants (APP1030243 to RLF and BAC; APP1107930) and Senior Research Fellowships (APP1079904, 606476), from the National Health and Medical Research Council of Australia, and the Outside Study Programme (Monash University). Generation of the *Nod1*^fl/fl^ mice was funded by the Public Health Agency of Canada. LY is supported by funding from the U. S. Department of Defense (Award No. W81XWH-17-1-0606). Research at the Hudson Institute of Medical Research is supported by the Victorian Government’s Operational Infrastructure Support Program.

## Author contributions

L. S. T. designed and performed experiments, analysed data and drafted sections of the manuscript. L. Y. performed experimental work, designed experiments and analysed data. K. D’C., G. W.-M., L. L., H. C., J. E., A. De P., M. K.-L. and J. C. performed experiments. G. K. designed and performed the organoid experiments. J. D., A. M., P. A. J., D. J. P., H. E. A. and U. N. provided expertise and essential research tools. S. C. and K. E. provided expertise and analysed imaging data. C. C. A., J. F. and T. A. K. generated important research tools. J. S. P. performed histological analyses. B. A. C. and S. L. M. designed and performed experiments. R. L. F. designed and performed experiments, analysed data, conceptualised the project and drafted the manuscript.

