## Supplementary material for "NOD1 mediates non-canonical inflammasome processing of interleukin-18 in epithelial cells to *Helicobacter pylori* infection": Supp. Fig. Legends

**Supplementary Figure Legends**

**Supplementary Fig. 1** *Il18*^-/-^ mice exhibit mucosal thickening of the stomach in response to chronic *Helicobacter* infection. **a-b** Images (**a**) and weights (**b**) of stomachs from *Il18*^+/+^ and *Il18*^-/-^ mice at 2 months p.i. with *H. pylori*. Arrows indicate thickening of the mucosa in an *Il18*^-/-^ animal. **c** Gastric cytokine and chemokine gene expression in the mice. **d-e** Images (**d**) and weights (**e**) of stomachs from *Il18*^+/+^ and *Il18*^-/-^ mice at 13 months p.i. with *H. felis*. Significance was determined by the unpaired two-tailed t-test. Data correspond to the mean ± SEM.

**Supplementary Fig. 2** BMDMs do not produce IL-18 in response to *H. pylori* stimulation. **a-b** IL-18 (**a**) and IL-1β (**b**) production in BMDMs from wild type (WT), *Nlrp3*^-/-^ and *Pycard*^-/-^ mice. BMDMs were either left untreated (UT) or stimulated with *H. pylori* bacteria (HP), *E. coli* LPS (LPS) alone or with the inflammasome activator, nigericin (LPS + Nig). Representative of two independent experiments. Significance was determined by the unpaired two-tailed t-test. Data correspond to the mean ± SEM.

**Supplementary Fig. 3** Mice lacking the canonical inflammasome proteins Nlrp1 and Nlrc4 are unaffected in *H. pylori* colonisation and pathology. **a-f** *Nlrp*^+/+^, *Nlrp1*^+/-^, *Nlrp*^-/-^ and **g, h** *Nlrc4*^+/+^ and *Nlrc4*^-/-^ mice were administered either BHI broth (control), *H. pylori* or *H. felis*. Mice were culled at 3 months (**a**-**f**) and 2 weeks (**g, h**) p.i. Stomachs from the mice were assessed for *H. pylori* bacterial loads (**a, h**), inflammation (**b-f**) and weights (**g**). Data pooled from two independent experiments (**g, h**).

**Supplementary Fig. 4** Human AGS gastric epithelial cells do not express detectable levels of the canonical inflammasome proteins, NLRP3 or ASC. **a-b** *NLRP3* (**a**) and *PYCARD* (**b**) expression in control AGS and *NOD1* KO CRISPR/Cas9 cell lines, as well as in control cell lines HeLa, HEK293T and THP-1. Cells were either left untreated (UT) or stimulated with *H. pylori* bacteria (HP). *RNA18S1* expression was determined as a loading control. **c** NLRP3 and ASC synthesis in cell lysates (LYS). Tubulin was used as a loading control.

**Supplementary Fig. 5** Nod1 deficiency has no effect on IL-18 production by BMDMs. **a-c** Total IL-18 (**a**), mature IL-18 (**c**) and IL-1β (**c**) production by BMDMs from *Nod1*^fl/fl^ and *Nod1*^fl/fl^ x *LysM*-cre mice. BMDMs were either left untreated (UT) or stimulated with *H. pylori* bacteria (HP), *E. coli* LPS (LPS) alone or with the inflammasome activator, nigericin (LPS+Nig). Data are representative of three independent experiments. Data correspond to the mean ± SEM.

**Supplementary Fig. 6** Ripk2 deficiency has no effect on IL-18 production. **a-b** Total IL-18 (**a**), mature IL-18 (**b**) production by GECs isolated from *Ripk2^+/+^* and *Ripk2^-/-^* mice. **c-d** Total IL-18 (**c**) and IL-1β (**d**) production by BMDMs isolated from *Ripk2^+/+^* and *Ripk2^-/-^* mice. BMDMs were either left untreated (UT) or stimulated with *H. pylori* bacteria (HP), HP outer membrane vesicles (MV), *E. coli* LPS (LPS) or LPS together with the inflammasome activator, nigericin. **e**, **f** Fold changes of Cxcl2 (**e**) and KC (**f**) production by HP-stimulated BMDMs from *Ripk2^+/+^* and *Ripk2^-/-^* mice normalised to the corresponding untreated cells. Data correspond to the mean ± SEM.

**Supplementary Fig. 7** Generation of conditional knockout mice lacking *Nod1*^-/-^ in the myeloid compartment. **a** C57BL/6 mice were generated in which the *Nod1* gene was “floxed” using a gene targeting strategy that involved deletion of *Nod1* exons 2 and 3, corresponding to the *Nod1* CARD. For this, a construct was generated in which *loxP* sites were situated external to these two exons and an FRT-flanked neomycin cassette used for selection. To generate mice lacking functional Nod1 in the myeloid compartment, *Nod1*^fl/fl^ mice were crossed with *lysM*-Cre animals. **b** *Nod1* PCR was performed on tail DNA and BMDM cDNA from two of these mice, using oligonucleotides specific for exons 2-3 and exon 7, respectively. DNA/cDNA samples were also tested from the progeny of *Nod1*^fl/fl^ mice crossed with epithelial cell-specific Cre (A33-Cre) animals and from *Nod1*^+/+^ mice crossed with *lysM*-Cre animals. Water was used as a negative control.
