## Supplementary figures and images for "NOD1 mediates non-canonical inflammasome processing of interleukin-18 in epithelial cells to *Helicobacter pylori* infection"

### Supp. Fig. 1

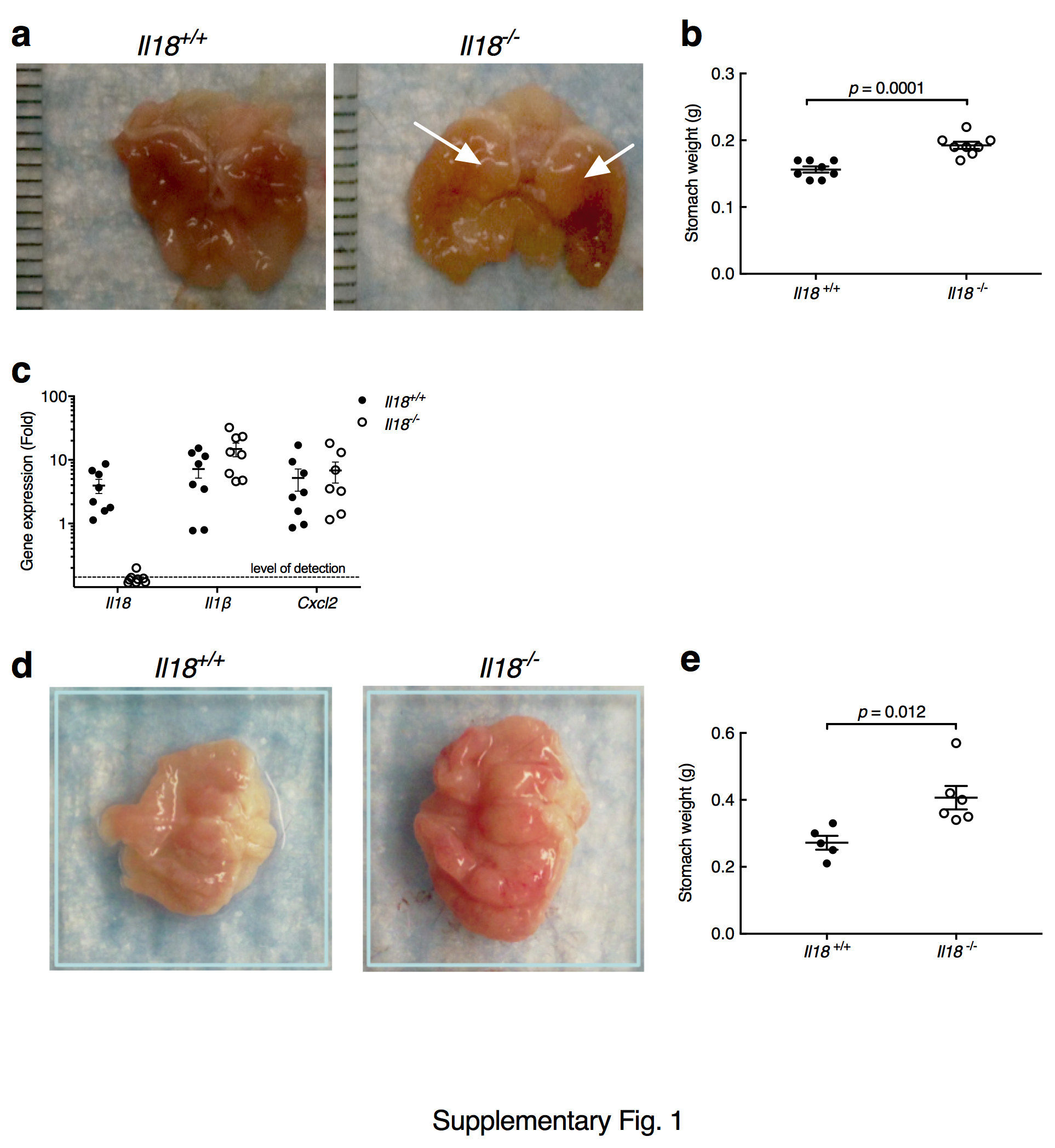

### Supp. Fig. 2

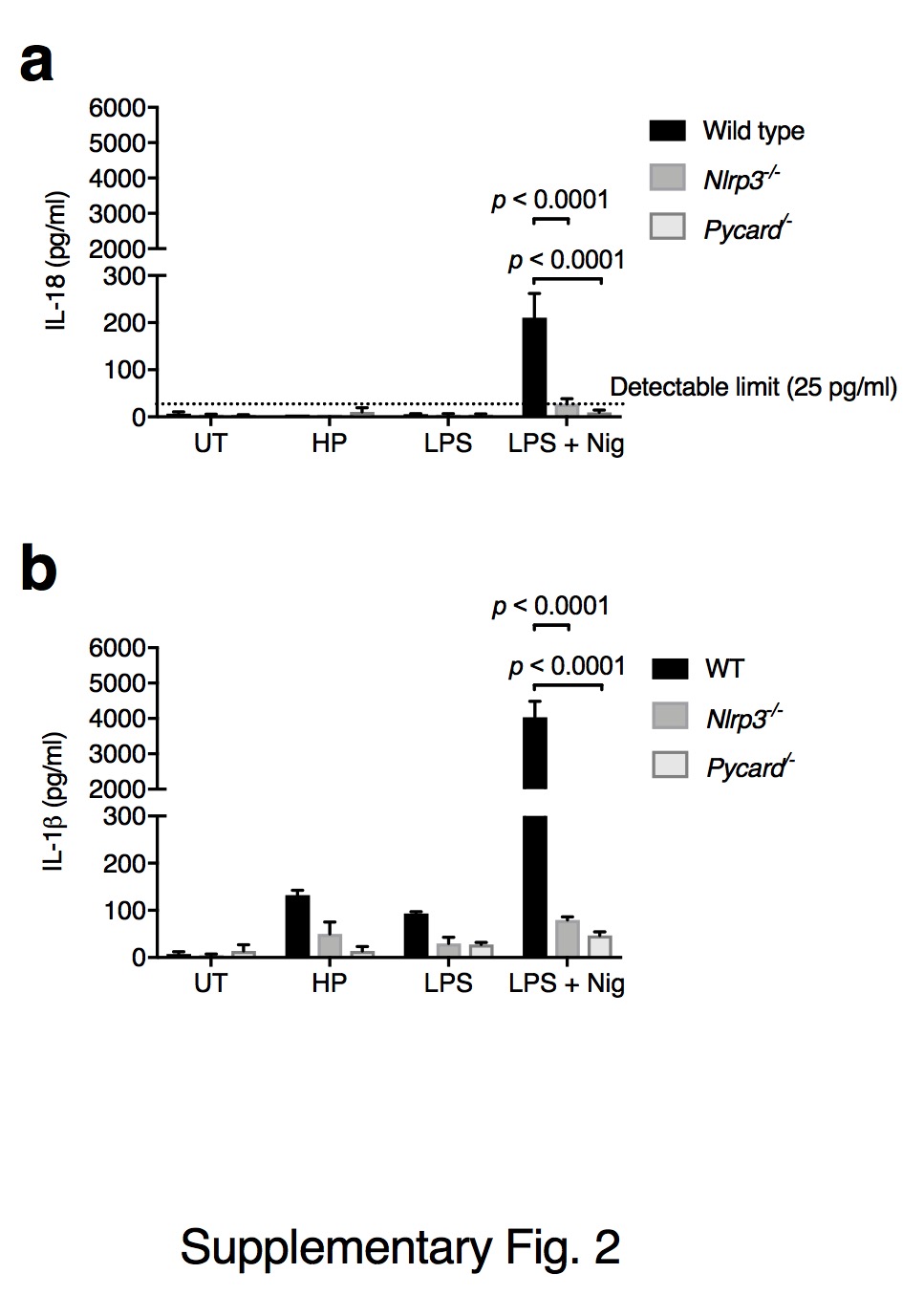

### Supp. Fig. 3

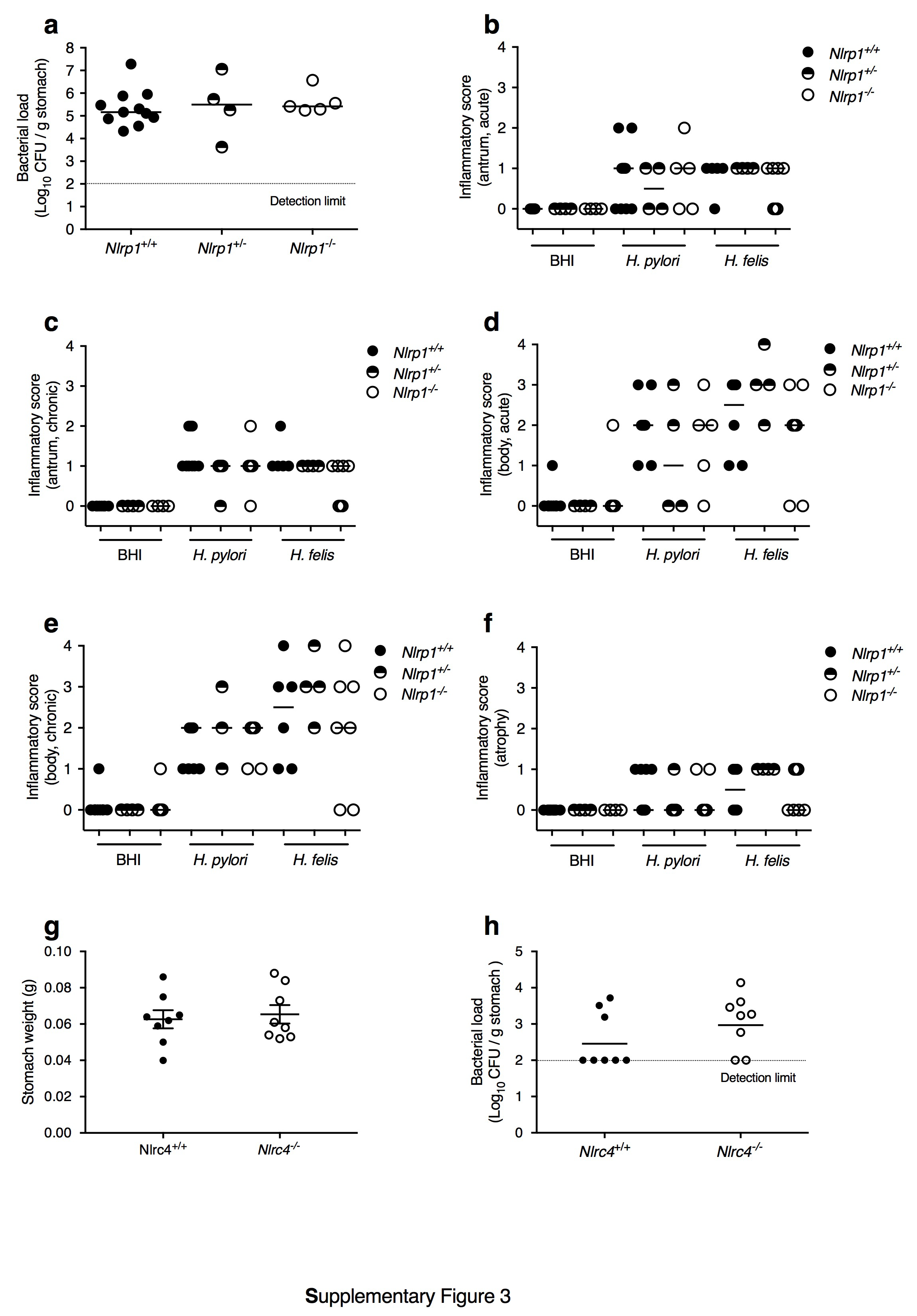

### Supp. Fig. 4

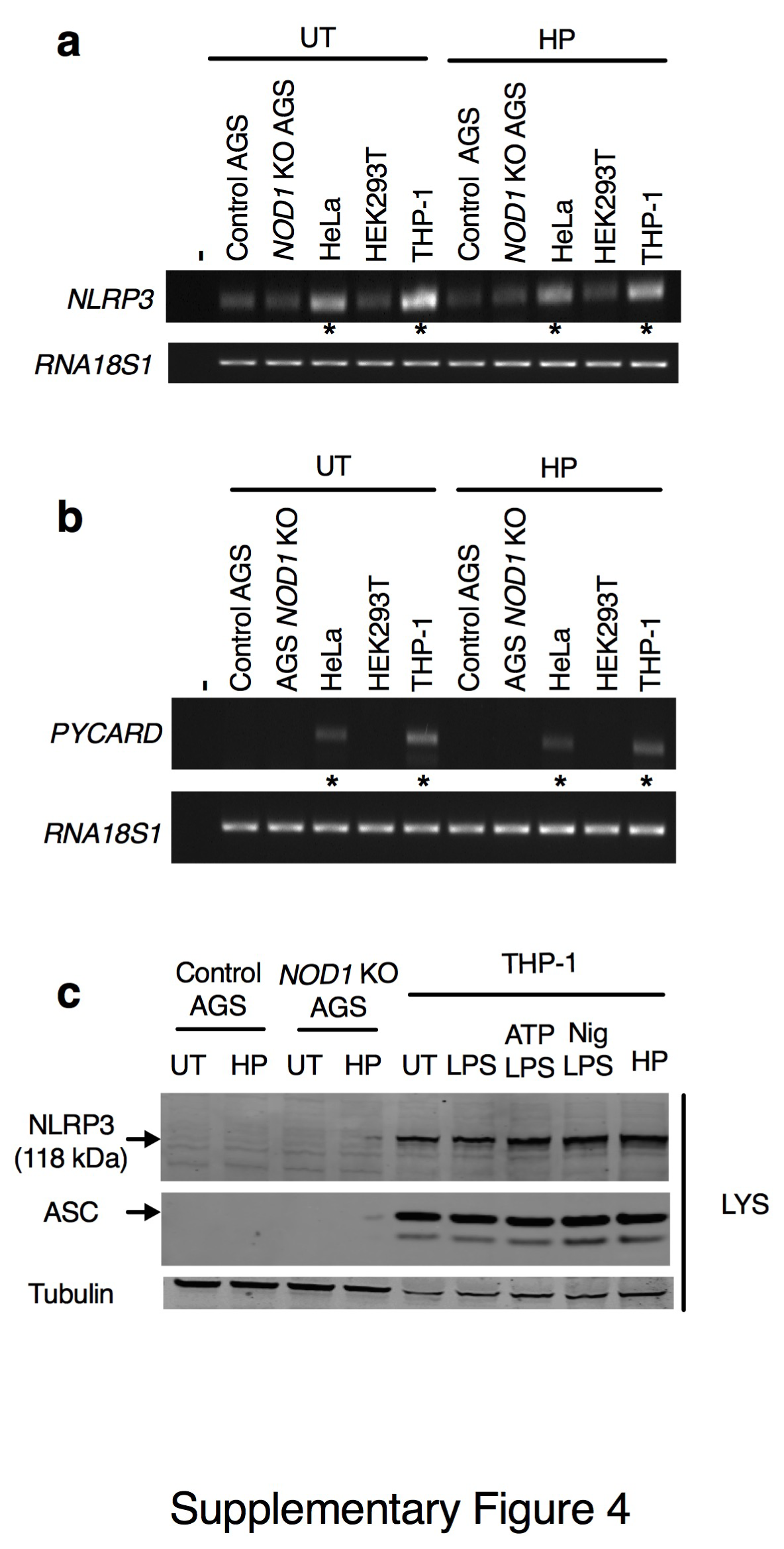

### Supp. Fig. 5

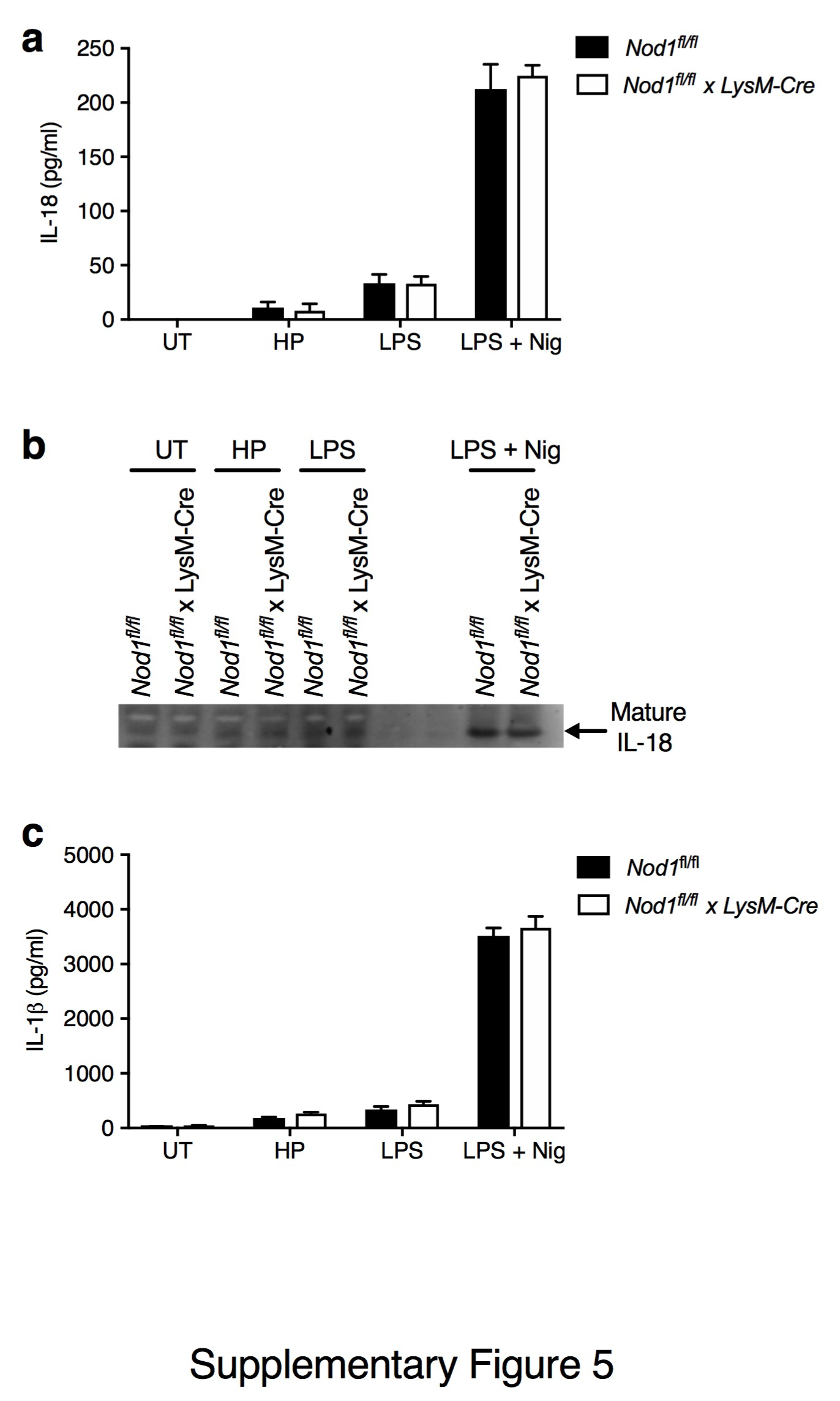

### Supp. Fig. 6

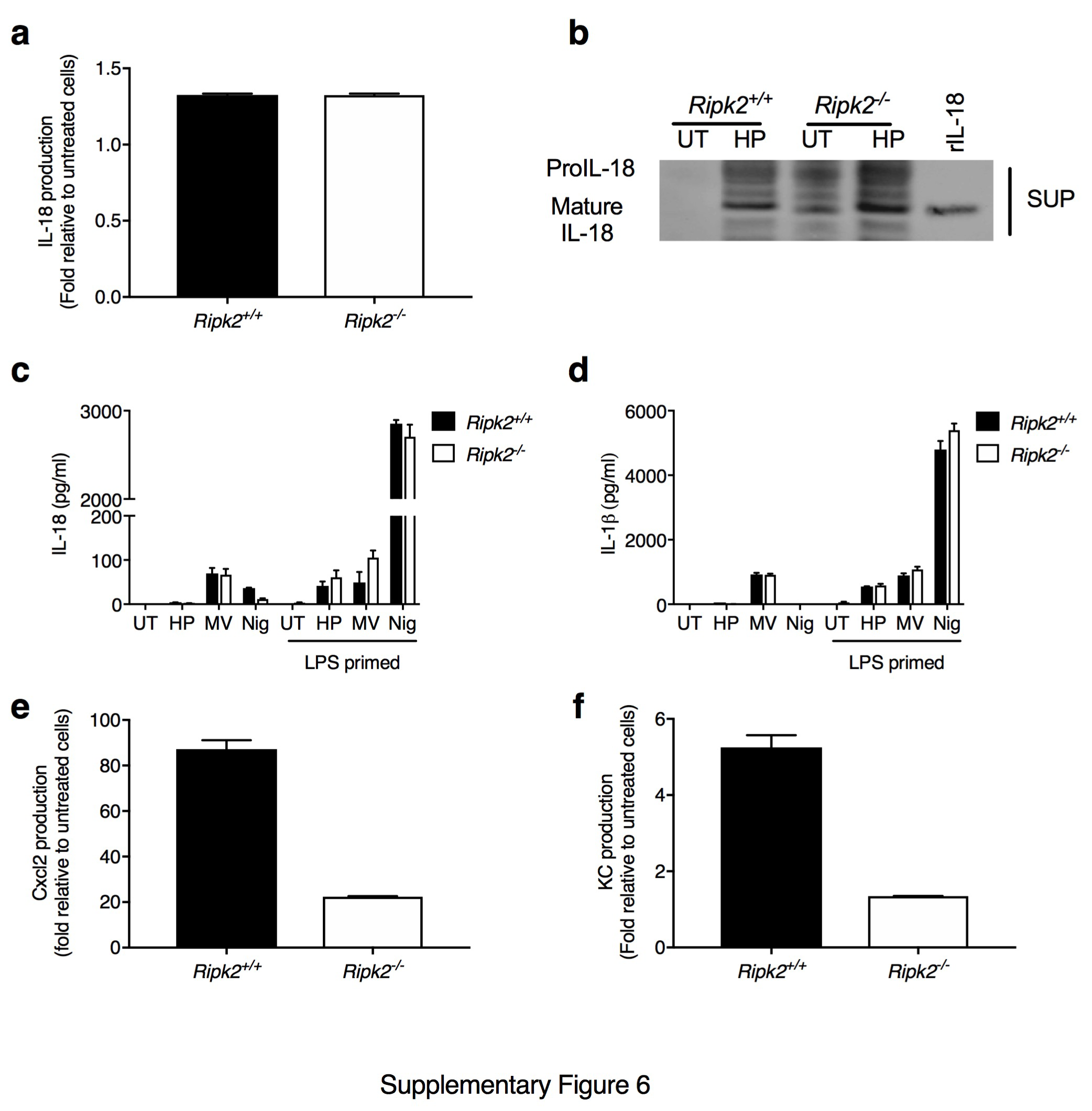
