## Supplementary material for "NOD1 mediates non-canonical inflammasome processing of interleukin-18 in epithelial cells to *Helicobacter pylori* infection": Supp. Fig. 7

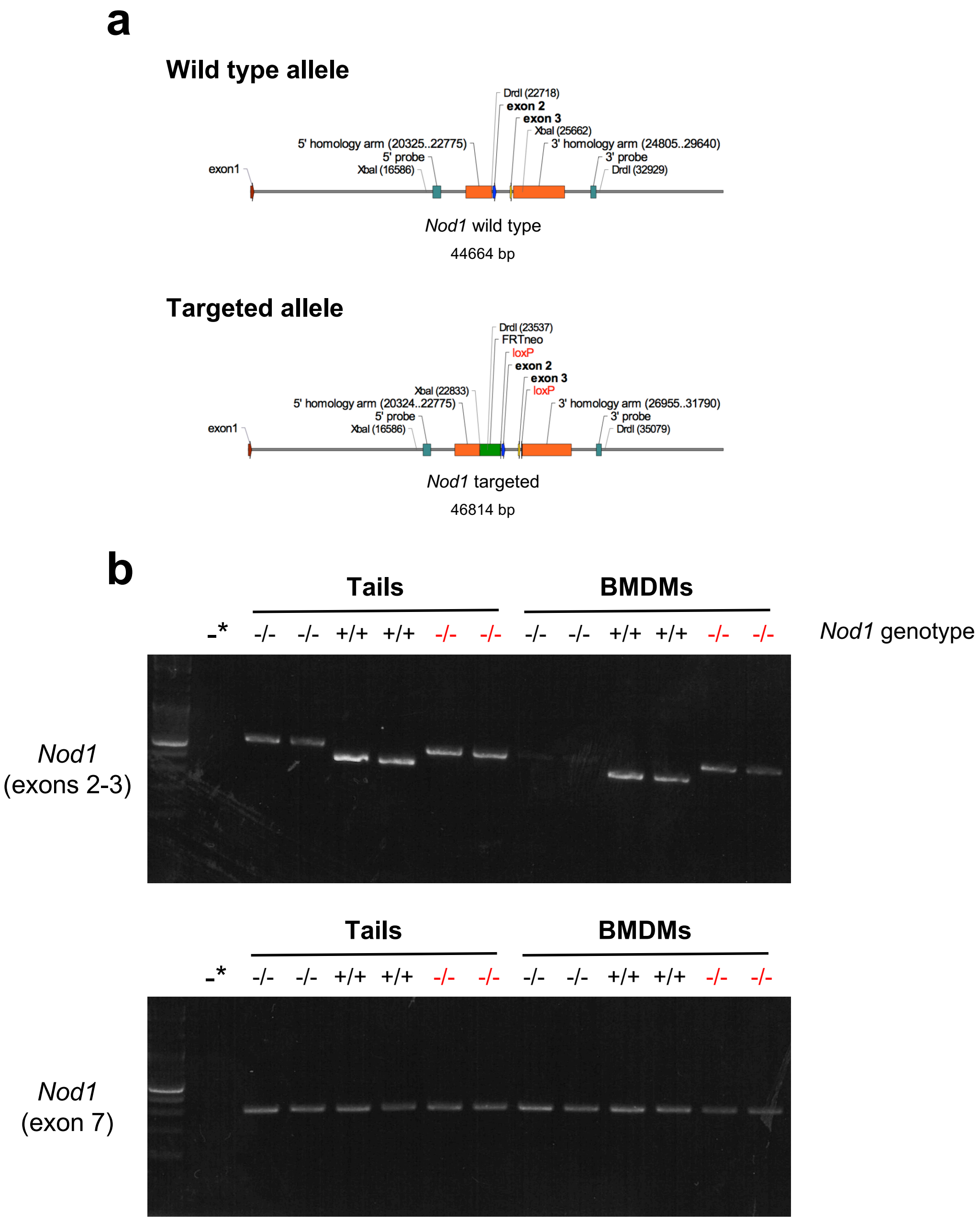


* water control

-/- = *Nod1*$^{fl/fl}$ x *LysM*-Cre (*Nod1* myeloid cell KO)
+/+ = *Nod1*$^{+/+}$ x *LysM*-Cre
-/- = *Nod1*$^{fl/fl}$ x A33-Cre (*Nod1* epithelial cell KO)

Supplementary Fig. 7
